# Predicting transdermal fentanyl delivery using physics-based simulations for tailored therapy

**DOI:** 10.1101/2021.01.21.427533

**Authors:** Flora Bahrami, René Michel Rossi, Thijs Defraeye

## Abstract

Transdermal fentanyl patches are an effective alternative to the sustained-release of oral morphine for chronic pain treatment. Due to the narrow therapeutic range of fentanyl, the fentanyl concentration in the blood needs to be controlled carefully. Only then, effective pain relief can be reached while avoiding adverse effects such as respiratory depression. In this study, a physics-based digital twin of the patient was developed by implementing mechanistic models for transdermal drug uptake and the patient’s pharmacokinetic and pharmacodynamics response. A digital twin is a virtual representation of the patient and the transdermal drug delivery system, which is linked to the real-world patient by patient feedback, sensor data of specific biomarkers, or customizing the twin to a particular patient characteristic, for example, based on the age. This digital twin can predict the transdermal drug delivery processes *in-silico*. Our twin is used first to predict conventional therapy’s effect for using fentanyl patches on a virtual patient at different ages. The results show that by aging, the maximum transdermal fentanyl flux and maximum concentration of fentanyl in the blood decrease by 11.4% and 7.0%, respectively. Nonetheless, by aging, the pain relief increases by 45.2% despite the lower concentration of fentanyl in the blood for older patients. As a next step, the digital twin was used to propose a tailored therapy, based on the patient’s age, to deliver fentanyl based on the patient’s needs to alleviate pain. This predesigned therapy consisted of customizing the duration of applying and changing the commercialized fentanyl patches based on the calculated pain intensity. According to this therapy, a patient of 20 years old needs to change the patch 2.1 times more frequently compared to conventional therapy, which led to 30% more pain relief and 315% more time without pain. In addition, the digital twin was updated by the patient’s pain intensity feedback. Such therapy led to an increase in the patient’s breathing rate while having effective pain relief, therefore providing a safer and more comfortable treatment for the patient. We quantified the added value of a patient’s physics-based digital twin and sketched the future roadmap for implementing such twin-assisted treatment into the clinics.

**Nomenclature:** *Symbols:* *c*_*i*_ The concentration of fentanyl in layer *i* (in the drug uptake model) [ng ml^-1^] *c*_*p*_ The concentration of fentanyl in the central compartment [ng ml^-1^] *c*_*r*_ The concentration of fentanyl in the rapid equilibrated compartment [ng ml^-1^] *c*_*s*_ The concentration of fentanyl in the slow equilibrated compartment [ng ml^-1^] *c*_*g*_ The concentration of fentanyl in the gastrointestinal compartment [ng ml^-1^] *c*_*l*_ The concentration of fentanyl in the hepatic compartment [ng ml^-1^] *c*_*e*_ The concentration of fentanyl in the effect compartment [ng ml^-1^] *D*_*i*_ Diffusion coefficient of fentanyl in layer *i* (in the mechanistic model) [m^2^ s^-1^] *D*_0_ Base diffusion coefficient of fentanyl [m^2^ s^-1^] *D*_*T*_ Diffusion coefficient of fentanyl at temperature T [m^2^ s^-1^] *D*_306_ Diffusion coefficient of fentanyl at 306[K] [m^2^ s^-1^] *d*_*pt*_ The thickness of the transdermal patch [µm] *d*_*sc*_ The thickness of the stratum corneum [µm] *d*_*vep*_ The thickness of the viable epidermis [µm] *d*_*Edm*_ The thickness of the equivalent dermis [µm] *E*_*i*_ The intensity of effect *i* 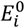 The baseline of effect *i* 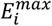 The maximum effect *i* *EC*_50,*i*_ The concentration related to half-maximum effect *i* [ng ml^-1^] *f*_*u*_ The fraction of unbound fentanyl in plasma *j*_*i*_ Fentanyl flux in layer *i* (in the mechanistic model) *K*_*i*/*j*_ The partition coefficient of fentanyl between layer *i* to *j* (in the mechanistic model) *K*_*i*_ The drug capacity in layer *i* (in the mechanistic model) *k*_*cs*_ Inter-compartmental first-order equilibrium rate constant (central to slow equilibrated) [min^-1^] *k*_*cr*_ Inter-compartmental first-order equilibrium rate constant (central to rapid equilibrated) [min^-1^] *k*_*cg*_ Inter-compartmental first-order equilibrium rate constant (central to gastrointestinal) [min^-1^] *k*_*ch*_ Inter-compartmental first-order equilibrium rate constant (central to hepatic) [min-1] *k*_*sc*_ Inter-compartmental first-order equilibrium rate constant (slow equilibrated to central) [min-1] *k*_*rc*_ Inter-compartmental first-order equilibrium rate constant (rapid equilibrated to central) [min^-1^] *k*_*hc*_ Inter-compartmental first-order equilibrium rate constant (hepatic to central) [min^-1^] *k*_*gh*_ Inter-compartmental first-order equilibrium rate constant (gastrointestinal to hepatic) [min^-1^] *k*_*met*_ Metabolization rate constant [min^-1^] *k*_*re*_ Renal clearance rate constant [min^-1^] *k*_*e*_ Inter-compartmental first-order equilibrium rate constant (for effect compartment) [min^-1^] *SI* Sensitivity index *t* Time [h] *t*_*D*_ Time lag [h] 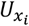 Dependent variable related to *x*_*i*_ for sensitivity analysis *V*_*c*_ The apparent volume of the central compartment [L] *V*_*s*_ The apparent volume of the slow equilibrated compartment [L] *V*_*r*_ The apparent volume of the rapid equilibrated compartment [L] *V*_*g*_ The apparent volume of the gastrointestinal compartment [L] *V*_*h*_ The apparent volume of the hepatic compartment [L] *x*_*i*_ The independent variable which sensitivity analysis is done based on it *γ* Hill coefficient *ψ*_*i*_ Drug potential in domain *i* [ng ml^-1^]

## 1 Introduction

Severe chronic pain is a frequent symptom in cancer patients, and about 70-80 % of the patients in advanced cancer stages deal with such pain [1]. However, proper pain treatment is challenging by considering the complex nature of pain and differentiated actions of opioids [2]. Untreated or poorly-treated pain could be overwhelming for the patients and will reduce their quality of life [3]. This pain causes some patients to fear pain even more than their disease, which may lead them to physician-assisted suicide [4]. Fentanyl is clinically used to treat moderate-to-severe cancer pain. Fentanyl is a synthetic opioid that is used in cases that non-steroidal anti-inflammatory drugs (NSAIDs) are insufficient [5]. As an alternative to oral and parenteral delivery, transdermal fentanyl delivery (TDD) is a clinically-approved therapy, which is successful due to the low molecular weight, high potency, and lipid solubility of fentanyl. The key advantage of transdermal fentanyl delivery besides simplicity and noninvasive delivery is that it offers a controlled delivery of fentanyl and avoids the first-pass metabolism [6], [7].

Transdermal fentanyl therapy shows high inter-individual and intra-individual variability [8]–[10]. Several factors can cause these variabilities [10], including a different clearance rate [11]. Additionally, fentanyl has a narrow therapeutic range. Besides the analgesic effect, fentanyl affects the ventilation rate in patients and could cause respiratory depression, which, in severe cases, could lead to patient death [12]. By considering these side effects, it is important to keep the fentanyl concentration in the blood at such a level that sufficient analgesia is reached while avoiding adverse effects such as respiratory depression for each patient. The conventional therapy for using the transdermal fentanyl patch, which is being used in clinics, is applying a transdermal patch for 72 hours on the skin [13], [14]. Applying a similar therapy for all the patients, by ignoring the physiological states and differences in the therapeutic range for each patient, often fails to treat the pain effectively [15]. Therefore, current practice involves a trial-and-error approach where the dose of the oral morphine is gradually increased while carefully monitoring the patient’s response until an individualized dosage and therapy are reached. With the advent of physics-based simulations of drug delivery processes, truly individualized therapies can be achieved [16].

Mechanistic, first-principles-based modeling and simulation of transdermal drug delivery systems are usually done using two approaches. The first approach is the simulation of the drug’s penetration process through the skin layers, especially the stratum corneum at different scales. In this approach, at the nano-scale, the penetration of drugs in lipid bilayers of the stratum corneum (SC) is modeled by using molecular dynamics simulation [17]–[19]. One of the results of this set of simulations is the diffusion coefficient for transient diffusion of the drug in the modeled layer [20], [21]. At the meso- and micro-scale, brick-and-mortar and cellular models of the SC are being used to model the drug transport [22]–[24]. Fickian diffusion-based models are typically used at the macro-scale to obtain drug penetration through the skin into the blood circulation system [25]–[27]. Such models can accurately capture the time-lag caused by the skin between the drug release and the drug uptake. In the second approach, pharmacokinetics (PK) and pharmacodynamics (PD) modeling and simulation are performed. In this approach, the skin is modeled as a depot to obtain the incoming flux of drugs to the blood circulation system [28], [29]. PK model calculates the concentration of drug in plasma, and the PD model provides the drug effect(s) related to the drug plasma concentration [30]–[34]. To propose a successful transdermal therapy *in-silico*, however, it is essential that both the delayed drug uptake kinetics through the skin are predicted *in-silico* together with the fate of the drug in the human body. Such a complete mechanistic model that follows the drug from the transdermal patch until the organ where it invokes pain relief has not been set up, to our best knowledge. In addition, by connecting the model to the real-world patient via patient physiology and pain feedback, a so-called digital twin of the patient for fentanyl delivery can be created to personalize therapy. Such a connection can be established by tailoring the digital twin to a certain patient or a certain class of patients (e.g., age group). More advanced digital twins are created by establishing real-time feedback of the patient to the twin, for example, the experienced pain scale or sensed biomarkers. Gartner predicted that by 2021, 50% of the large industrial companies would rely on digital twins, leading to an expected 10% gain in effectiveness [35], [36]. Implementing digital twin in healthcare enables *in silico* trials on a population of virtual patients [37], and there were studies on digital twin in training surgeon through interactive virtual simulations [38], or digital twin for aerosol pulmonary drug therapy [39], [40]. However, the digital twin concept is still novel, and its full potential has not been exploited so far, including for transdermal drug delivery.

In this paper, we quantify the added value of such physics-based digital twins for effective pain management for patients with different age classes as a step toward tailored fentanyl transdermal therapy. By coupling the drug uptake model, PK, and PD models, we developed a tailored digital twin for a virtual patient at different ages. The PK model was validated, the PD model was calibrated based on the literature, and the uptake model was validated in our previous study [41]. By using these digital twins, in a first step, we studied the effect of age on the outcome of the same fentanyl transdermal therapy. In the second step, we controlled the incoming flux of drugs to the body for the virtual patient at different ages in a way to have sufficient pain relief while avoiding hypoventilation. Finally, the digital twin was connected to the virtual patient by real-time feedback to modify the predesigned therapy based on the virtual patient’s need.

## 2 MATERIALS AND METHODS

### 2.1 Mechanistic multiphysics model for transdermal drug delivery

The developed physics-based digital twin in this study consisted of 3 model blocks to simulate transdermal drug uptake, PK, and PD models. In the transdermal drug uptake model, the flux of fentanyl from the patch through the skin into the blood circulation system was calculated. The PK model calculated fentanyl distribution through the body, metabolization in the liver, and elimination by the kidneys. The result of the PK model is the plasma fentanyl concentration versus time. In the PD model, the fentanyl concentration in the effect compartment was calculated. Based on this concentration, pain relief and ventilation rate as the effects of fentanyl were obtained. The overall structure of this physics-based digital twin is shown in Figure 1.

**Figure 1.**
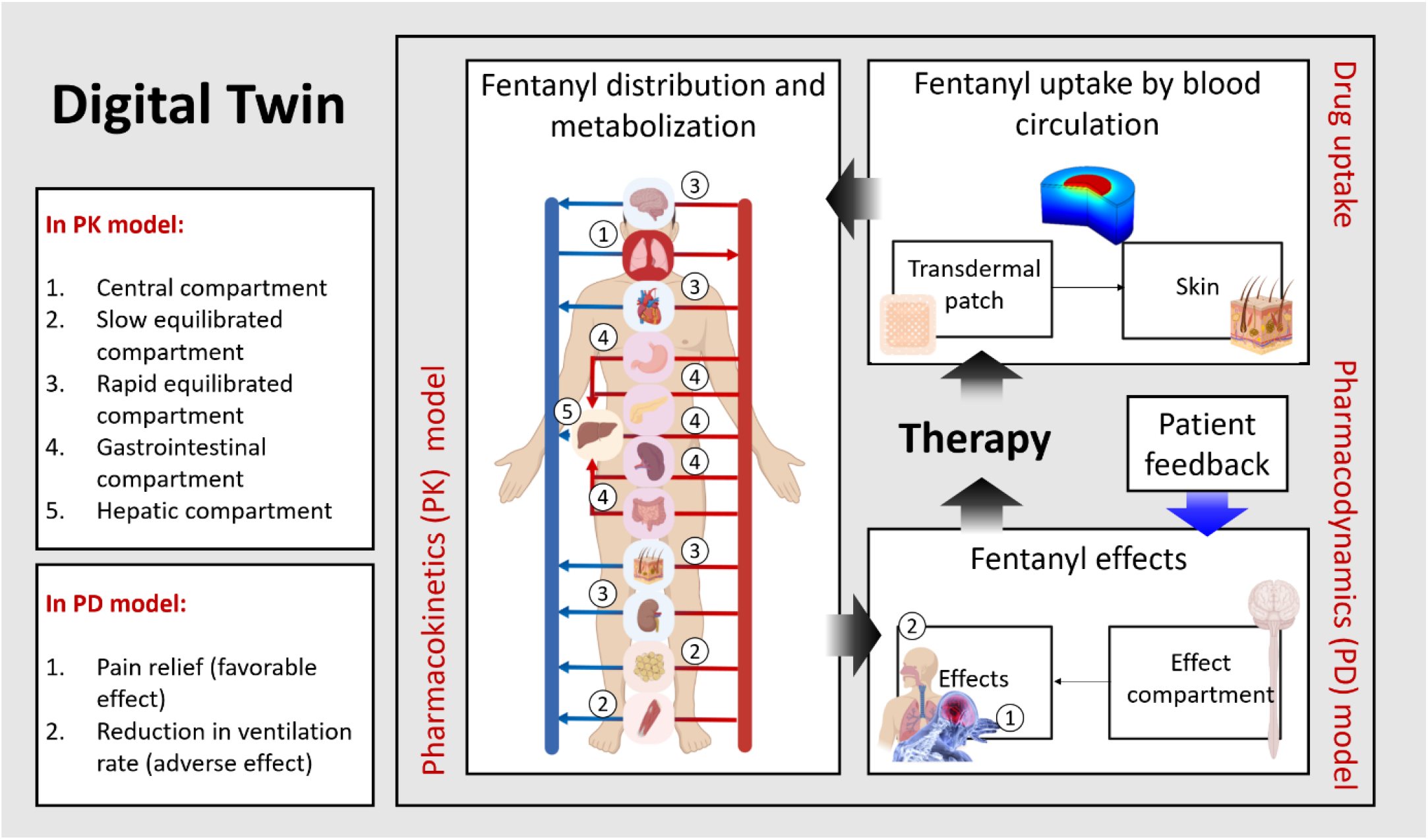
Overall structure of an implemented digital twin for transdermal fentanyl therapy (Created with BioRender.com).

#### 2.1.1 Transdermal drug uptake in skin

##### 2.1.1.1 Computational system configuration

A mechanistic continuum model was built in order to simulate fentanyl release from a transdermal patch, diffusion through the skin, and uptake by blood circulation. Based on fentanyl properties, such as high lipophilicity and low molecular weight, fentanyl is suitable to be used via the first generation of transdermal patches. In a way, the penetration of fentanyl through the skin barrier is diffusion-driven [6], and there is no need for penetration enhancers. The model and simulation were built and executed according to best practice guidelines (or clinical practice guidelines) in modeling for medical device design [42], [43].

The geometry of the drug uptake model and boundary conditions are shown in Figure 2. The system configuration consists of four blocks with a square surface area representing fentanyl patch, stratum corneum, viable epidermis, and a part of the dermis. Transdermal patches are designed to deliver the drug at an approximately constant rate [44]. Therefore, the fentanyl patches are commercially labeled with a targeted drug release rate of 12-100 µg h-1, which is the average rate over 72 h [7].

**Figure 2.**
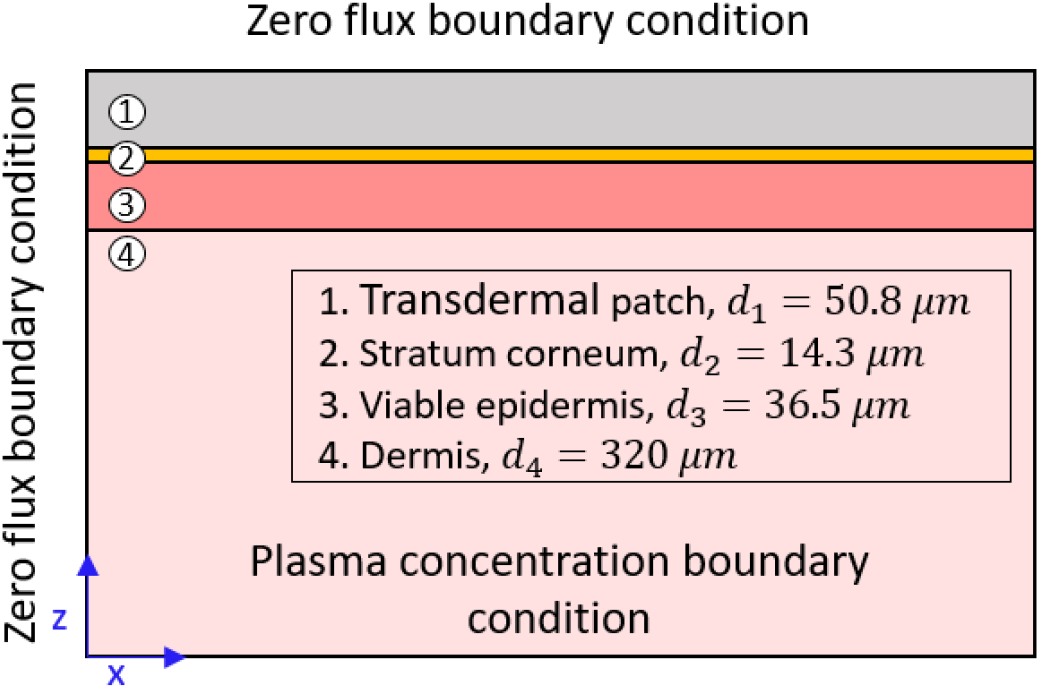
Geometrical model of the transdermal patch and skin layers.

We chose the patch’s size based on a Duragesic® fentanyl patch with a nominal flux of 75 µg h-1, which contains 7.5 mg of fentanyl [13]. The patch’s surface area is 30 cm2, and the thickness of the patch was considered 50 µm. Here, we used the diffusion parameters for the transdermal patch and epidermis layer (stratum corneum and viable epidermis) from our previous study [41], [45]. In that study, the model parameters’ values were optimized to reach the best match with experimental data, where the epidermis was considered a single layer. The value of these parameters is mentioned in section 2.1.1.3, Table 1.

**Table 1.**
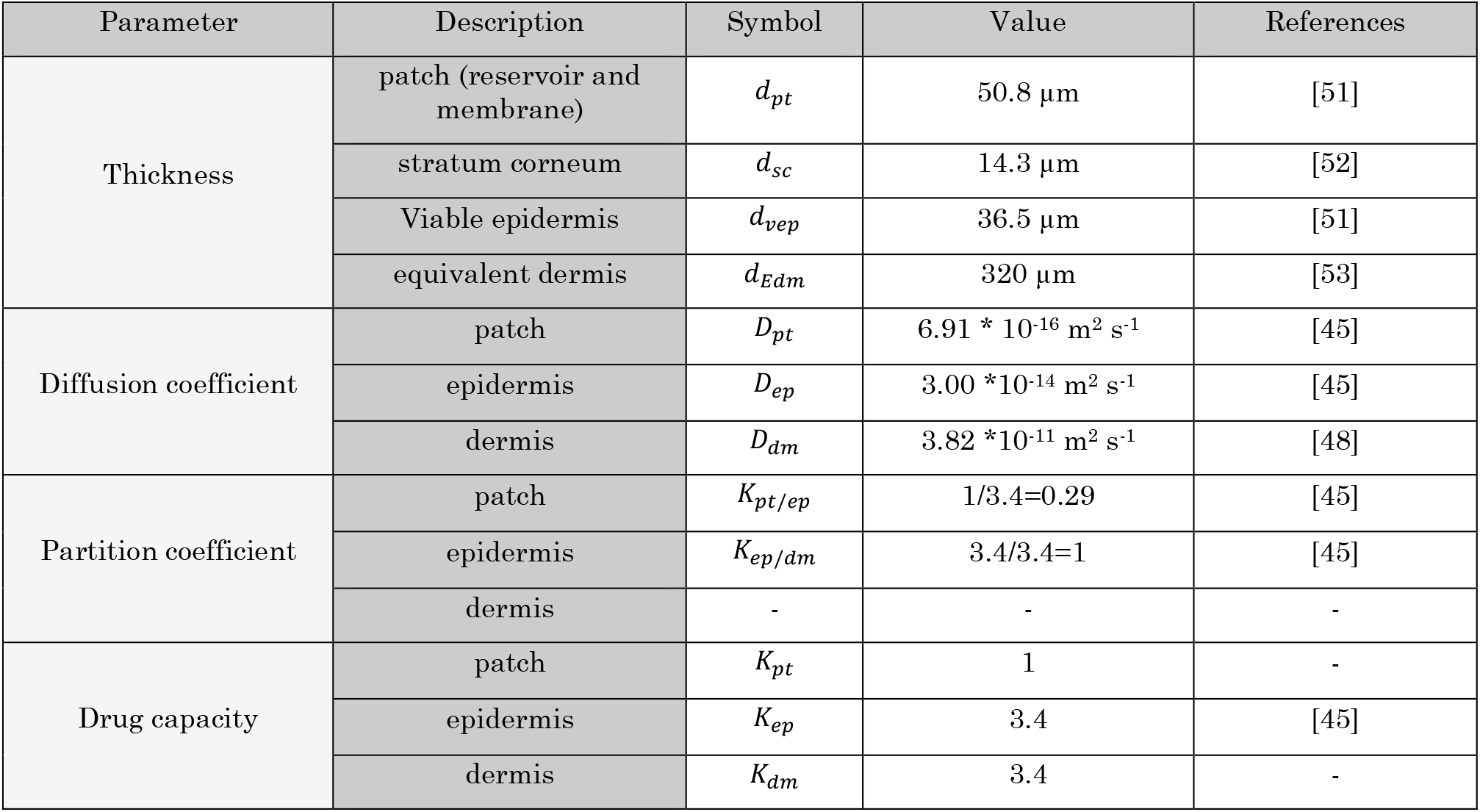
Parameters involved in the mechanistic model for 20 years old virtual patient.

| Parameter | Description | Symbol | Value | References |
| --- | --- | --- | --- | --- |
| Thickness | patch (reservoir and membrane) | $d_{pt}$ | $50.8 \mu\text{m}$ | [51] |
| | stratum corneum | $d_{sc}$ | $14.3 \mu\text{m}$ | [52] |
| | Viable epidermis | $d_{vep}$ | $36.5 \mu\text{m}$ | [51] |
| | equivalent dermis | $d_{Edm}$ | $320 \mu\text{m}$ | [53] |
| Diffusion coefficient | patch | $D_{pt}$ | $6.91 * 10^{-16} \text{ m}^2 \text{ s}^{-1}$ | [45] |
| | epidermis | $D_{ep}$ | $3.00 * 10^{-14} \text{ m}^2 \text{ s}^{-1}$ | [45] |
| | dermis | $D_{dm}$ | $3.82 * 10^{-11} \text{ m}^2 \text{ s}^{-1}$ | [48] |
| Partition coefficient | patch | $K_{pt/ep}$ | $1/3.4=0.29$ | [45] |
| | epidermis | $K_{ep/dm}$ | $3.4/3.4=1$ | [45] |
|  | dermis | - | - | - |
| Drug capacity | patch | $K_{pt}$ | 1 | - |
| | epidermis | $K_{ep}$ | 3.4 | [45] |
| | dermis | $K_{dm}$ | 3.4 | - |

Due to the presence of capillary vessels in the dermis, the drug would be uptaken by the vessels after the penetration of the drug from the epidermis. These vessels are in different depths in the dermis, and the drug uptake happens in various lengths of the dermis. In this study, we assumed that the drug was transferred through the dermis only by the diffusion process, which is a simplified assumption. If we consider the whole length of the dermis by only applying the diffusion process, the time lag increases up to 11 h (refer to Equation 5, section 2.1.1.2.), which is considerably different from reported values (1 to 2 h time-lag [46]). In the real situation, due to plasma flow into and out of the vessels, there is an advection mechanism as well. As a result, the drug penetration mechanism in the dermis is a combination of diffusion and advection. However, to modify this assumption, instead of modeling the dermis’s whole thickness, we considered an equivalent diffusive length of the dermis. The dermis’s equivalent length was obtained based on the reported experimental data of time-lag for detecting fentanyl in the plasma [46] and the diffusion coefficient of fentanyl in the dermis.

##### 2.1.1.2 Governing equations

The transport of fentanyl in the patch and skin was modeled by one-dimensional transient diffusion. The fentanyl concentration in the patch and skin layers is denoted by *c*_*i*_ (*z, t*) [kg m^-3^]. This diffusion process is described by Fick’s second law, which is mentioned in Equation 1 [26].

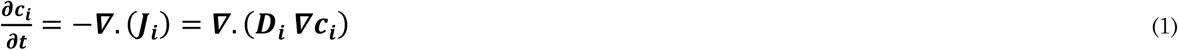

Where *J*_*i*_ and *D*_*i*_ are the flux [kg m^-2^ s^-1^] and diffusion coefficient [m^2^ s^-1^] of fentanyl in domain *i*, respectively. At the interface of two layers (patch and skin layers), due to the different solubility of drugs in different layers, partitioning should be considered. The partition coefficient at the interface is defined in Equation 2 [26].

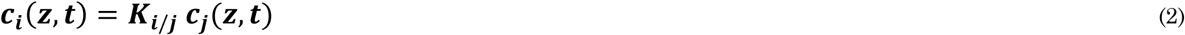

Here, *K*_*i*/*j*_ is the partition coefficient of fentanyl from domain *i* to domain *j*. As a result of partitioning, a discontinuity will occur at the interface, which is inconvenient to solve computationally. To avoid this problem, another variable was defined, which is continuous throughout the whole domain. The relation of this new variable *ψ*_*i*_ and *c*_*i*_ is mentioned in Equation 3.

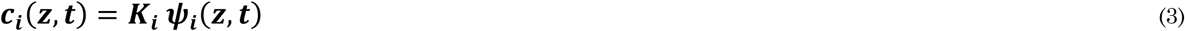

In this equation, *K*_*i*_ and *ψ*_*i*_ are drug capacity and drug potential in domain *i*. As mentioned before, we assumed that *ψ*_*i*_ is continuous; therefore, at the interface, the drug potential in domain *i* and *j* are equal. By applying this condition in Equations 2 and 3, we arrive at Equation 4.

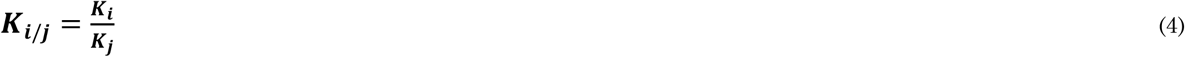

The equivalent length of the dermis was considered based on the time lag of detection fentanyl after applying the patch on the skin. Between the application of the first fentanyl patch (100 or 75 µg h^-1^) and detection of fentanyl in blood circulation (> 0.1 ng ml^-1^), there is 1 to 2 h time-lag [46]. Based on the reported range, we considered the time-lag as 1.5 [h]. Both epidermis and dermis are playing a role in this delay. We calculated the time-lag caused by the epidermis by using the mechanistic model. After subsiding the delay caused by the epidermis, the dermis was responsible for the remaining part. By using Equation 5, the equivalent diffusive length of the dermis was obtained.

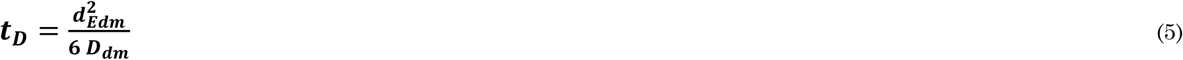

In this equation, *t*_*D*_ is the time lag [s] and *d*_*Edm*_ [m] is the equivalent length of the dermis [47]. The diffusion coefficient of fentanyl in the dermis is shown by *D*_*dm*_ [m^2^ s^-1^], which obtained by Equation 6.

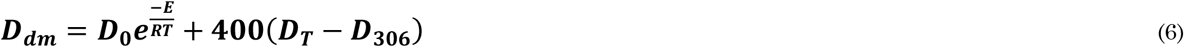

Which, *D*_0_ is the base diffusion coefficient, *D*_306_ is the diffusion coefficient at *T* = 306 K, and *D*_*T*_ is the diffusion coefficient in operating temperature, which is 310.5 K [48].

##### 2.1.1.3 Material properties and transport characteristics of skin and patch

The drug used in this simulation is fentanyl with a molecular weight of 336.5 g mol-1. The material properties used in this study are mentioned in Table 1. In this study, we assumed the fentanyl capacity of the dermis to be the same as the epidermis. The Log P (octanol-water partition coefficient) of fentanyl is reported between 1.5 to 4.3 [49], and based on the study that has been done by Cross et al. 2003; there is a correlation between Log P and the partition coefficient between epidermis and dermis layer [50]. By considering the range of log P for fentanyl, the partition coefficient between epidermis and dermis is between 0.4 −9, which our assumption lies in the range.

##### 2.1.1.4 Boundary and initial conditions

In this mechanistic model, the source of fentanyl is the transdermal patch, which has an initial concentration of 80 kg m^-3^, equivalent to 12.6 mg of fentanyl. This amount of drug is based on the Duragesic® fentanyl patch with 75 µg h^-1^ flux of drug. As the initial condition, it is assumed that there is no fentanyl in the skin layers. We assumed that there is no flux in peripheral boundaries, and the flux of the drug is vertical from the patch to the end of the dermis. At the bottom boundary of the dermis, the concentration is equal to the fentanyl concentration in blood circulation, which is zero at the beginning (the concentration of fentanyl in blood circulation is obtained by the PK model).

#### 2.1.2 Physiologically-based pharmacokinetics modeling

The physiologically based PK model was used to calculate the concentration of fentanyl in plasma by considering drug metabolization, elimination, and distribution through the body. The PK model was developed based on the physiology of the patient and the pharmacological mechanism of fentanyl. In PK modeling, it is common to lump different organs into a single functioning kinetic unit, despite the fact that they are anatomically distinct (Upton et al., 2016). To capture better the complexity of fentanyl transport in the human body and the resulting drug concentration in the blood plasma, five different kinetic compartments were used in the present study:

1. A central compartment, namely the blood circulation system and lungs (Equation 7).
2. A slowly equilibrating compartment, which includes muscles, carcass, and fat tissue (Equation 8).
3. A rapidly equilibrating compartment, which includes the brain, heart, skin, and kidneys (Equation 9)
4. A gastrointestinal compartment, which includes the spleen, gut, and pancreas. In this compartment, the outcoming flow goes to the hepatic compartment instead of the central compartment (Equation 10).
5. A hepatic compartment, where the metabolization of fentanyl is occurring. This is the main elimination route of fentanyl from the human body (Equation 11).

An overview of the PK model is given in Figure 1. To evaluate the concentration of fentanyl in each compartment, the law of conservation of mass by considering well-mixed compartments were applied. By using the first-order kinetic rate law, the fentanyl concentration at each compartment based on transferring drug between compartments, metabolization, and elimination was calculated. The ordinary differential equations related to mass conservation for each compartment are mentioned below:

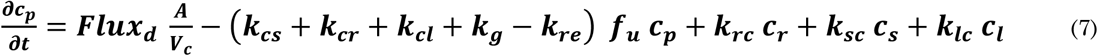

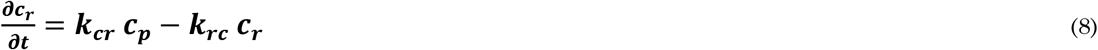

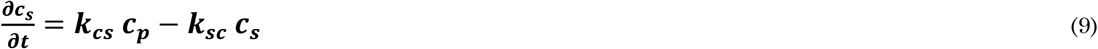

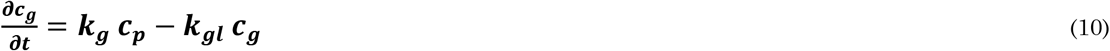

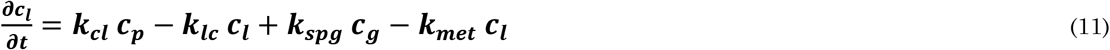

In this set of equations, *c*_*p*_, *c*_*r*_, *c*_*s*_, *c*_*g*_, and *c*_*l*_ are concentration of fentanyl in central, rapid equilibrated, slow equilibrated, gastrointestinal and hepatic compartments, respectively. *f*_*u*_ is the unbound fraction of fentanyl in the central compartment. *k*_*ij*_, *k*_*met*_, and *k*_*re*_ are the first-order equilibrium rate constants for inter-compartmental clearance, metabolization, and renal clearance, respectively. The parameters used in these equations are specified in Table 2.

**Table 2.** Parameters involved in the pharmacokinetics model.

| Parameter | Description | Symbol | Value | Reference |
| --- | --- | --- | --- | --- |
| The apparent volume of the compartment (for fentanyl) | central compartment | $V_c$ | 23.8 L | [30] |
| | slow equilibrated compartment | $V_s$ | 808 L | |
| | rapid equilibrated compartment | $V_r$ | 24.5 L | |
| | gastrointestinal compartment | $V_g$ | 17.3 L | |
| | hepatic compartment | $V_l$ | 20.4 L | |
| Inter-compartmental first-order equilibrium rate constant | central to slow equilibrated | $k_{cs}$ | $1.5 \cdot 10^{-3} \text{ s}^{-1}$ | [30], [55] |
| | central to rapid equilibrated | $k_{cr}$ | $1.2 \cdot 10^{-3} \text{ s}^{-1}$ | |
| | central to gastrointestinal | $k_{cg}$ | $5 \cdot 10^{-4} \text{ s}^{-1}$ | |
| | central to hepatic | $k_{ch}$ | $1.2 \cdot 10^{-4} \text{ s}^{-1}$ | |
| | slow equilibrated to central | $k_{sc}$ | $5 \cdot 10^{-5} \text{ s}^{-1}$ | |
| | rapid equilibrated to central | $k_{rc}$ | $1.2 \cdot 10^{-3} \text{ s}^{-1}$ | |
| | hepatic to central | $k_{hc}$ | $7 \cdot 10^{-4} \text{ s}^{-1}$ | |
| | gastrointestinal to hepatic | $k_{gl}$ | $5 \cdot 10^{-5} \text{ s}^{-1}$ | |
| Elimination of constant rate | metabolization | $k_{met}$ | $4.2 \cdot 10^{-3} \text{ s}^{-1}$ | [56] |
| | renal clearance | $k_{re}$ | $3 \cdot 10^{-5} \text{ s}^{-1}$ | |
| The unbound fraction of fentanyl | central compartment | $f_u$ | 0.22 | [57] |

#### 2.1.3 Mechanism-based pharmacodynamics model

In the PD model, the effects of fentanyl - pain relief and reduction of ventilation rate - were calculated. In fentanyl pharmacology, the drug effect will lag behind the fentanyl concentration in the plasma. As fentanyl rapidly binds to µ-receptors in the central nervous system (CNS), the biophase distribution model can approximate the fentanyl concentration in the effect compartment based on plasma concentration [58]. The effect compartment is a theoretical concept that is assumed to have the same distributional properties as the drug’s site of action [59]. This effect compartment is used to describe the time course between plasma drug concentration and the effect. [60]. The biophase concentration of fentanyl was obtained by using a standard effect-compartment equation (Equation 12).

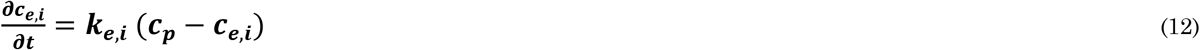

Where *c*_*e,i*_ is the fentanyl concentration in the effect compartment, *k*_*e,i*_ is the first-order equilibrium rate constant, and *i* represents the effect (subscript *VAS* for pain relief and subscript *rd* for reduction in ventilation rate). The values for parameters of PD model are brought in Table 3.

**Table 3.** Parameters involved in the pharmacodynamics model for 20 years old virtual patient.

| Effect | Parameter | Symbol | Value | Reference |
| --- | --- | --- | --- | --- |
| Ventilation rate | first-order equilibrium rate constant | $k_{e,rd}$ | $7 * 10^{-4} \text{ s}^{-1}$ | [58] |
| | baseline effect | $E_{rd}^0$ | $20 \text{ L min}^{-1}$ | |
| | maximum effect | $E_{rd}^{max}$ | $12 * 0.91 \text{ L min}^{-1}$ | |
| | Half maximal effective concentration | $EC_{50,rd}$ | $1.14 \text{ ng ml}^{-1}$ | |
| | hill coefficient | $\gamma_{rd}$ | 2.68 | |
| VAS pain score | first-order equilibrium rate constant | $k_{e,VAS}^{**}$ | $2 * 10^{-3} \text{ s}^{-1}$ | [56] |
| | baseline effect | $E_{VAS}^0$ | 7 | - |
| | maximum effect | $E_{VAS}^{max}$ | 7 | - |
| | Half maximal effective concentration | $EC_{50,VAS}$ | $1.35 \text{ ng ml}^{-1}$ | [62] |
| | hill coefficient | $\gamma_{VAS}$ | 2.7 | |

**\*\*This value is for analgesic effect**

##### 2.1.3.1 Drug effect

Fentanyl molecules target opioid receptors, which are mainly located in the brain within specialized neuroanatomical structures, which control emotion, pain, and addictive properties. Activation of opioid receptors by fentanyl will produce analgesia [61]. On the other hand, fentanyl activates opioid receptors on neurons within respiratory networks of the brainstem, which might lead to opioid-induced respiratory depression [60]. However, respiratory depression does not occur for all patients under fentanyl transdermal therapy. Here we track the ventilation rate as a factor to evaluate the risk of respiratory depression. In this study, the changes in ventilation rate (related to respiratory depression) and VAS pain score (to measure pain relief) were considered the main effects of fentanyl on the patient. A sigmoidal model based on the maximum effect (*E*_*max*_) best describes the fentanyl effects related to pain relief and reduction of ventilation rate. Maximum effect (*E*_*max*_) represents the maximum possible deviation from baseline effect. The baseline effect is the physiology state before using the drug, which in this case the baseline effects are initial pain intensity and initial ventilation rate. The intensity of pharmacologic effects of fentanyl for pain relief and ventilation rate based on maximum effect, and Equation 13 calculates fentanyl concentration in effect compartments.

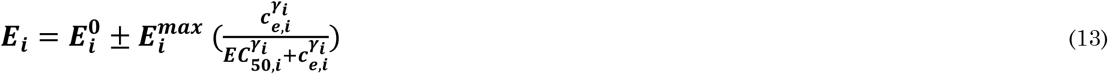

In Equation-13, 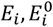, and 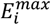 are the intensity of the pharmacologic, baseline, and maximum reachable effect, respectively. Here, the pharmacologic effect is the intensity of pain or minute ventilation of the patient. The baseline effect shows the state of pain or the patient’s minute ventilation before the administration of the drug. *EC*_50,*i*_ is the concentration of drug in plasma which corresponds to half of the achievable effect of the drug. In this study, *EC*_50,*VAS*_ (for pain relief) is the concentration when the pain intensity reaches half of the initial pain intensity and *EC*_50,*rd*_ (for the reduction in ventilation rate) is the concentration when the ventilation rate reaches half of its initial rate, and γ_*i*_ is the Hill coefficient [58].

The PD model parameters related to the reduction of ventilation rate were obtained from values reported in the literature [58]. The values used for these parameters are mentioned in Table 3. The corresponding values of the PD model for pain relief were obtained by fitting a sigmoidal model to experimental data from the literature [62], and the calculated values are brought in Table 2. In modeling the VAS pain score, we assumed that the initial VAS pain score for all the virtual patients is equal to 7, which lies in the range of severe pain. Another assumption is that we considered the cause of the pain is constant throughout the therapy. This assumption implies that if we remove the patch after the concentration of fentanyl reaches zero, the VAS pain score will return to its initial level.

#### 2.1.4 Validation and calibration of the models

The validation of the models was done separately for all models. The skin model’s validation for transdermal drug uptake is described elsewhere [41], [41].

##### 2.1.4.1 Validation of the PK model

To validate the pharmacokinetics model, we used the experimental data of the concentration of fentanyl in the plasma over time from Marier et al. (2006). In this series of experiments, they analyzed 24 men aged 18-45 years, with a body mass of at least 60 kg and a body mass index of 18-26 kg/m^2^. The participants were medically healthy, with clinically normal electrocardiograms (ECGs), and with no history of alcohol and drug abuse. The analysis was done over 11 days, with three fentanyl commercial patches with the nominal flux of 50 µg h^-1^, each for 72 hours. The PK parameters, which were used in this validation, are mentioned in Table 2, and the parameters for the mechanistic model, which indicates the incoming flux of fentanyl, are mentioned in Table 1. The result of this validation is brought in the supplementary material. The result of the validation of the PK model is mentioned in supplementary materials.

##### 2.1.4.2 Calibration of the PD model for pain relief

To develop a PD effect model for the VAS pain score, we used the reported VAS pain scores via fentanyl plasma concentration of Sandler et al. (1992) work. In this study, 29 adult patients, aged between 18 to 80, and with weight less than 100 kg, undertreatment of intravenous fentanyl with an infusion rate of 1.0 µg kg^-1^ h^-1^ were analyzed. The model for the VAS pain score was assumed sigmoidal, which is mentioned in Equation 20. The baseline effect (*E*_0,*VAS*_), maximum effect (*E*_*max,VAS*_, which here it is equal to *E*_0,*VAS*_), concentration of the half-maximum effect 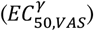, and Hill coefficient (*γ*) were obtained via fitting the model to experimental data, and the resulting RMSD was 0.471. The result of the calibration of the PD model is mentioned in supplementary materials.

### 2.2 Spatial and temporal discretization

The grid for the finite element method in each layer of the skin and patch was built based on the grid sensitivity analysis. In this sensitivity analysis, the spatial discretization error was considered 0.1% of the outcoming flux from the dermis layer, based on Richardson extrapolation. Therefore, the entire system’s grid, including transdermal patch and skin layers, consisted of 110 quadrilateral finite elements. The accumulation of grids is higher near the interface area to increase numerical accuracy. The duration of the simulations was 9 days (216 hours), which represents three periods of using fentanyl transdermal patch based on accepted therapy in the clinics. Based on the sensitivity analysis, the maximum time step should be 6 h. However, in this series of simulations, smaller time steps (0.1 h) were chosen to get a higher temporal resolution of the result.

### 2.3 Numerical implementation and simulation

COMSOL Multiphysics® software (version 5.5, COMSOL AB, Stockholm, Sweden), a finite-element-based commercialized software, was used in this study. The code developers verified this software. Therefore, the authors did not perform additional code verification. The mass transfer process of fentanyl through the patch and skin (drug uptake model) was carried out by the finite element method with partial differential equations interface (coefficient form). Quadratic Lagrange elements were used with a fully coupled direct solver, which relied on the MUltifrontal Massively Parallel sparse direct Solver (MUMPS) solver scheme. The drug distribution in body, metabolization, and elimination (PK model) was modeled using the ordinary differential equations interface (in boundary domain). To track the changes in the concentration of fentanyl in the effect compartment (PD model), an ordinary differential equations interface was used. For evaluating the effect of fentanyl, based on the effect compartment fentanyl concentration at each time step, the boundary probe was implemented. The boundary probe was calculating the effect of the drug at each time step based on the current concentration of the drug in the effect compartment. The tolerances for solver settings and convergence were determined by means of sensitivity analysis in such a way that a further increase in the tolerance did not alter the resulting solution.

### 2.4 Computational configurations

#### 2.4.1 Conventional fentanyl transdermal therapy

In clinics, the conventional therapy for fentanyl is to use a fentanyl patch for 72 hours and replace the patch, and the new patch needs to be placed on a new location body to avoid possible skin irritation [13]. To consider a period of therapy that includes the effect of changing patches in the process, we chose the duration of applying three patches (9 days). In our model, as mentioned earlier, we changed the new patch’s location; however, we assumed that the skin properties remain unchanged for different skin locations.

#### 2.4.2 Effect of age on the model parameters

The thickness of each layer of the skin changes by increasing the age. The effect of age on each layer is not similar, for instance, as age increases, the thickness of SC will increase, while the thickness of the dermis will decrease (Boireau-Adamezyk et al. 2014; Robert and Robert 2009). It should be noted that the changes in the dermis layer based on age were modified in this study to predict the changes in the equivalent thickness of the dermis based on age. Equations 14 [52] and 15 [53] show the age’s effect on each layer’s thickness.

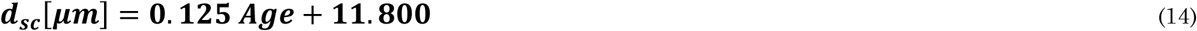

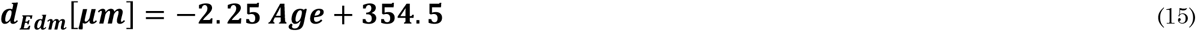

The PK parameters do not show significant changes based on age [65]; therefore, in this study, we assume the PK parameters are the same for the virtual patient of different ages. Despite PK parameters, Scott and Stanski (1987) showed that the required opioid dose for reducing the intensity of electroencephalography (EEG) reduces by increasing age. They studied the effect of fentanyl on 20 patients with age between 20 to 88 years. They were all male and without any renal and hepatic disease. They found a relationship between age and concentration of the half-maximum effect of fentanyl for the effect of the reduction of EEG. In this study, we assumed the normalized rate of changes in EC_50_ for the reduction of EEG by age is similar to the reduction of VAS pain score. The derived equation for changes in half-maximal effective concentration for VAS pain score is mentioned in Equation 16.

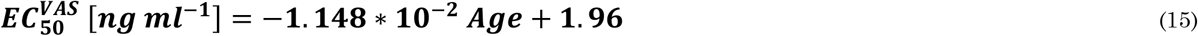

#### 2.4.3 Individualized therapy with a precalibrated digital twin

In the first step, we did not consider the virtual patient feedback in the model, which we refer to this twin as a precalibrated digital twin. The twin is customized to the patient class by calibrating it in advance for a certain age class, using equations 14, 15, and 16. By using this precalibrated digital twin, the VAS pain score of the virtual patient was calculated at each time. Based on the pain level of the virtual patient, the decision of keeping the patch on or changing it will be made. In this process, the type of fentanyl patch has not been changed, and it has the properties of a Duragesic® fentanyl patch with a nominal flux of 75 µg h^-1^. The considered criteria for changing the patch are that the VAS pain score is above 3, while the gradient of pain is positive (pain is increasing). A VAS pain score at or under 3 means the pain level is in the mild range; therefore, we considered the VAS pain score 3 as the target treatment to check for changing the patch. When the VAS pain score goes under 3, the patch will remain on the skin until the VAS pain score goes above 3, again. The strategy of changing the patch based on the pain is represented in a flowchart in Figure 4/a.

**Figure 3.**
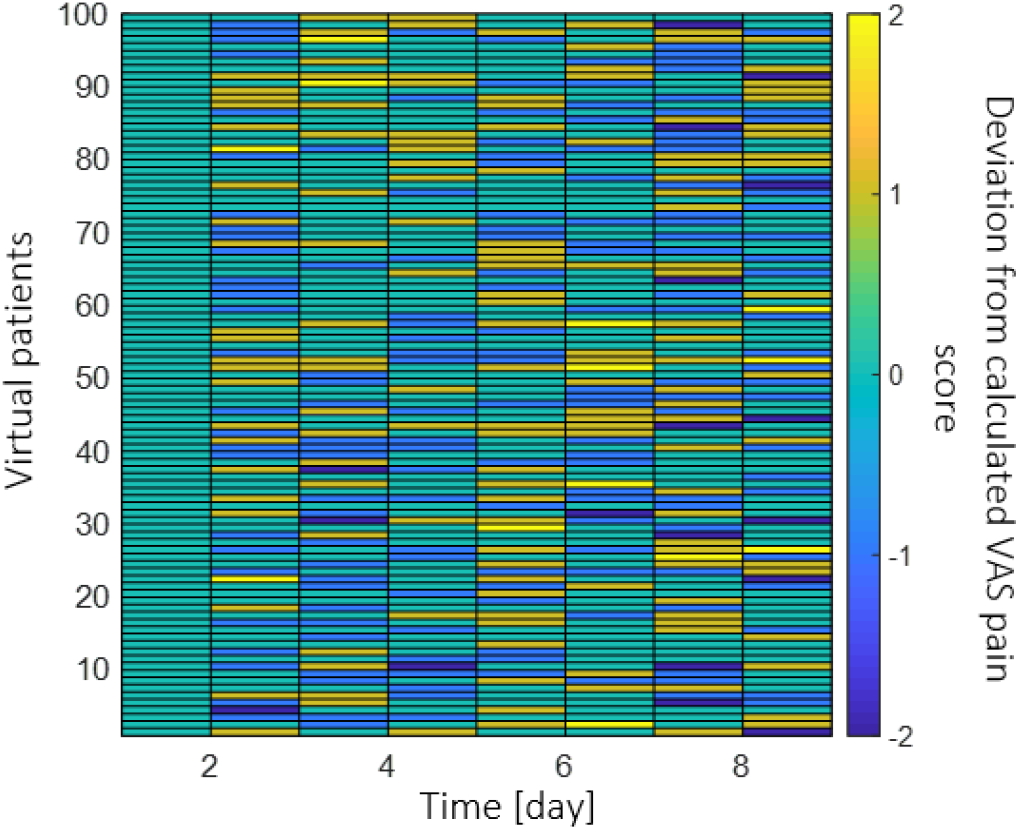
The set of random deviation from the calculated VAS pain score for 100 virtual patients at 8 feedback moments for 9 days.

**Figure 4.**
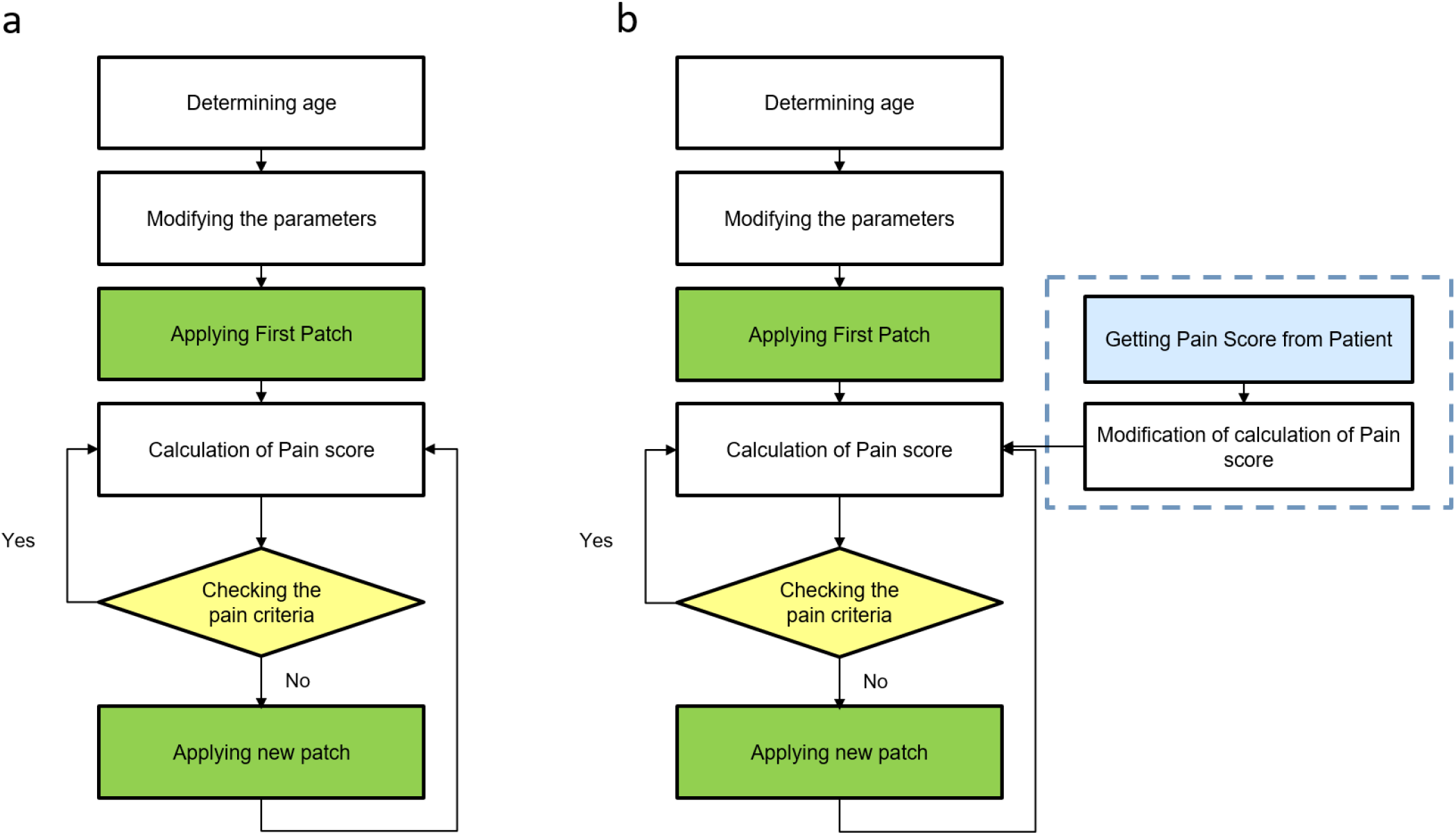
a: Flowchart of making the decision for changing the patch based on the VAS pain score calculation. b: - Flowchart of making the decision for changing patch based on VAS pain calculation updated by virtual patient feedback.

#### 2.4.4 Personalized therapy real-time digital twin

Due to the complex nature of the human body and the changes that happen to patient response, the calculated VAS pain score might have a deviation from the real VAS pain score. To take this effect into account, we mimicked a different patient response for 100 patients. With a real-time digital twin, the patient’s VAS pain score will be taken from the virtual patient at defined time intervals, and the calculation will be updated with these data. To include the virtual patient’s real VAS pain score in the model, we assumed that the virtual patient would enter the VAS pain score (integer numbers between 0-10) every 24 hours to the model. Another assumption for patient feedback was that the maximum difference between patient pain intensity and calculated pain is two VAS units. We produced a set of random feedback by considering the mentioned assumptions (deviation from calculated pain by PD model) via MATLAB®. The produces random deviation from calculated pain is shown in Figure 3. The flowchart of the strategy of updating the digital twin based on the virtual patient VAS pains score data is shown in a flowchart in Figure 4/b.

### 2.5 Metrics

The process of drug delivery from the patch to the site of action was analyzed by several metrics. These metrics are remaining drug in the patch (m_pt_), total delivered drug to the skin (m_ts_), and amount of drug in the skin (m_s_) as a function of time. Another important metric in this study is time without pain, which was considered the time that the VAS pain score is under 3 during the treatment period.

The remaining drug content (m_pt_ (t)) in the patch is being calculated as a function of time, which being reduced based on outgoing flux from the patch. The calculation for mpt (t) is brought in Equation 17.

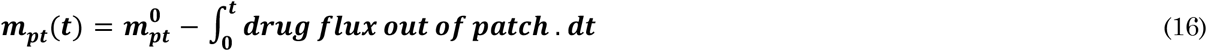

Where m^0^_pt_ is the initial amount of drug in the patch. The total delivered drug (m_ts_ (t)) to the body can be calculated based on the patch’s remaining drug. Equation 18 demonstrates the calculation of the total delivered drug to the body.

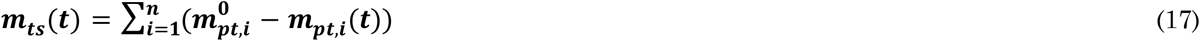

Where n is the number of patches that are used during a certain period of therapy. The amount of drug in the skin is related to the incoming flux of drug from the patch and outgoing flux to the blood circulation. This variable can be calculated based on the concentration of drug in skin layers which being calculated by Fick’s second law. The calculation of the amount of drug in the skin (ms(t)) is shown in Equation 19.

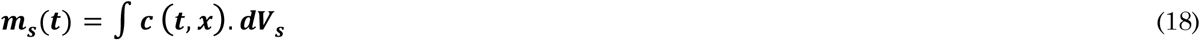

Where V_s_ is the volume of the skin, regarding the time without pain, as previously mentioned, we sum up the times that the VAS pain score is under 3.

### 2.6 Sensitivity analysis of the model parameters

The sensitivity of the total delivered drug from the skin (m_ts_), average fentanyl plasma concentration during 72 hours 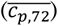 and the average VAS pain score to the related parameters were studied. These parameters are related to drug uptake (partition coefficient, diffusion coefficient, layer’s thickness), PK (compartment’s volume, blood flow between compartments, clearance coefficient, and the fraction of unbound drug), and PD (half-maximal effective concentration and Hill coefficient) model. The sensitivity index in this study was calculated for each parameter based on Equation 20.

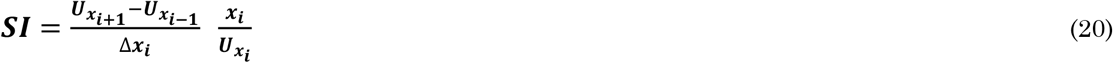

Where x_i_ is the model input parameter and U_xi_ is the dependent variable (m_ts_, 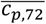 or VAS pain score) corresponded to x_i_. SI is the sensitivity index, which we use to analyze the sensitivity of U_xi_ to xi. Here, we considered 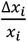 as 1%, to evaluate the sensitivity index based on this 1% deviation from the chosen value for xi. The result of sensitivity analysis is shown in supplementary materials.

## 3 RESULTS and DISCUSSION

### 3.1 Analysis of conventional fentanyl transdermal therapy

At first, the drug uptake, PK, and PD simulations were performed for conventional transdermal therapy for a virtual patient at age 20 years. In Figure 5, the fentanyl flux released from the patch and out of the dermis is shown. When a patch has been placed, there is a peak in the drug flux out of the patch. The maximal flux out of the patch for the first patch was 398 µg cm^-2 h-1^, which occurred in one hour of applying the patch. After one hour of applying the patch, the outgoing flux of fentanyl from the patch decreases drastically; in a way, the outgoing flux of fentanyl reaches 6.85 µg cm^-2 h-1^ after 1 h and 0.95 µg cm^-2 h-1^ after 72 h. After releasing the drug from patch to skin, due to the low diffusion coefficient and thickness of skin, there is a time lag to reach the maximum flux of outgoing drug from the skin. As the drug gradually leave the dermis, the outgoing drug flux from the skin does not change drastically, unlike the patch. As shown in Figure 8, the maximal flux out of the dermis was 2.31 µg cm^-2 h-1^, and it occurred after 19 hours after applying the patch, and the average flux out of the dermis during 72 hours is 1.47 µg cm^-2 h-1^. The delay in drug uptake between the dermis and patch is clear and spans several hours. This shows the slow response and thus the need for proper control of transdermal therapy.

**Figure 5.**
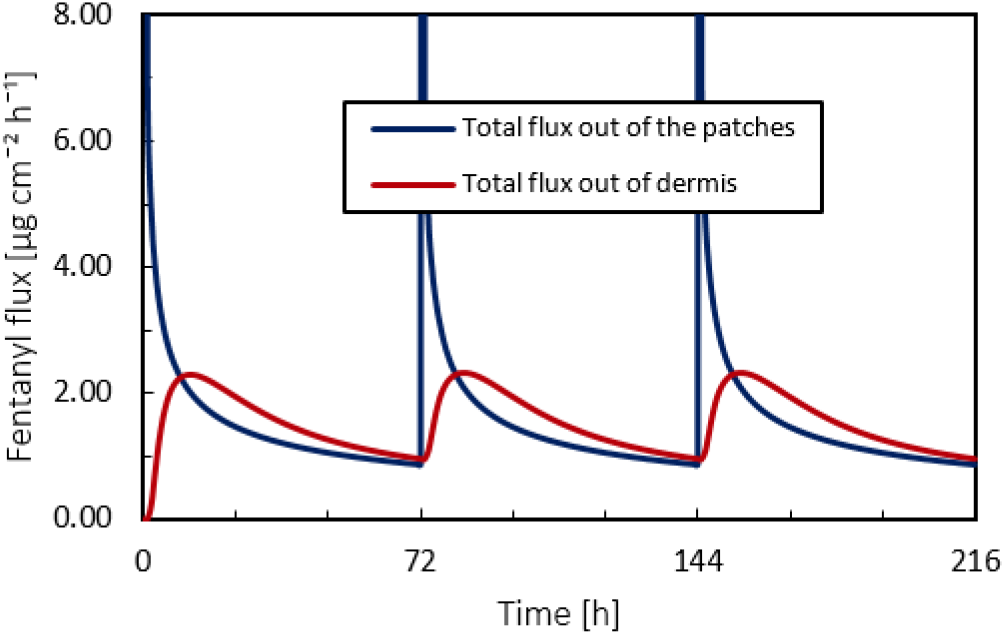
Fentanyl flux at skin-patch and dermis-blood circulation interface during 9 days by applying 3 patches of fentanyl with the nominal flux of 75 µg h-1 each for 72 hours.

**Figure 6.**
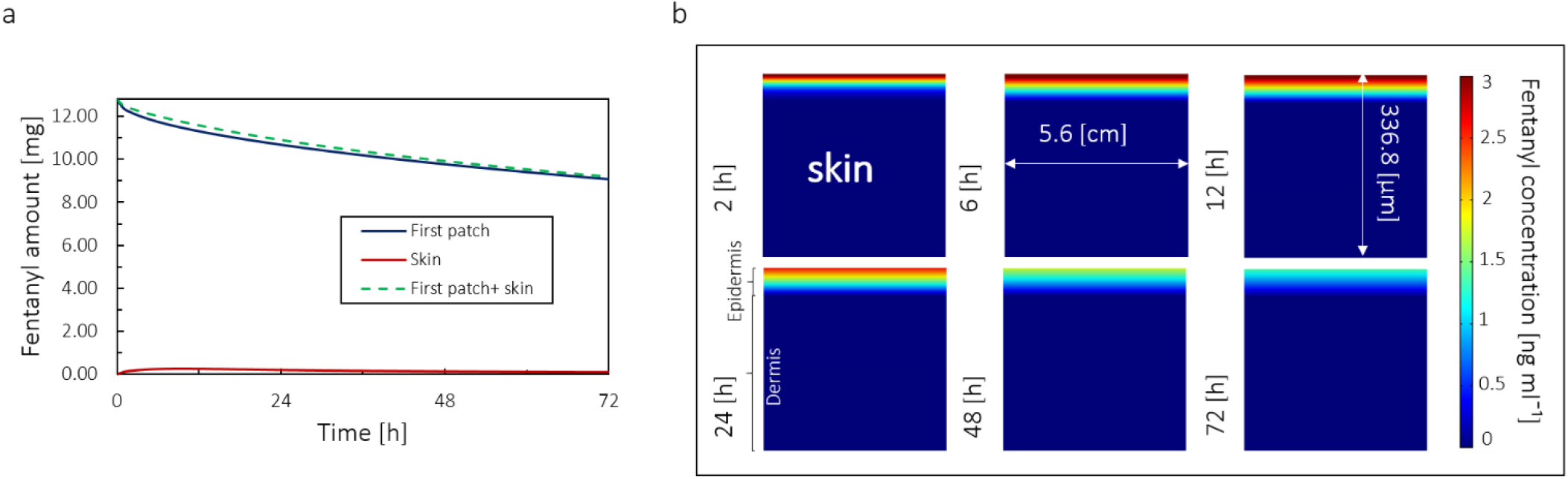
a: Amount of fentanyl stored in the patch, skin, and cumulative amount fentanyl in skin and patch in 72 h by applying one patch. b: Concentration of fentanyl in skin layers in 72 h for the virtual patient at the age of 20 years.

**Figure 7.**
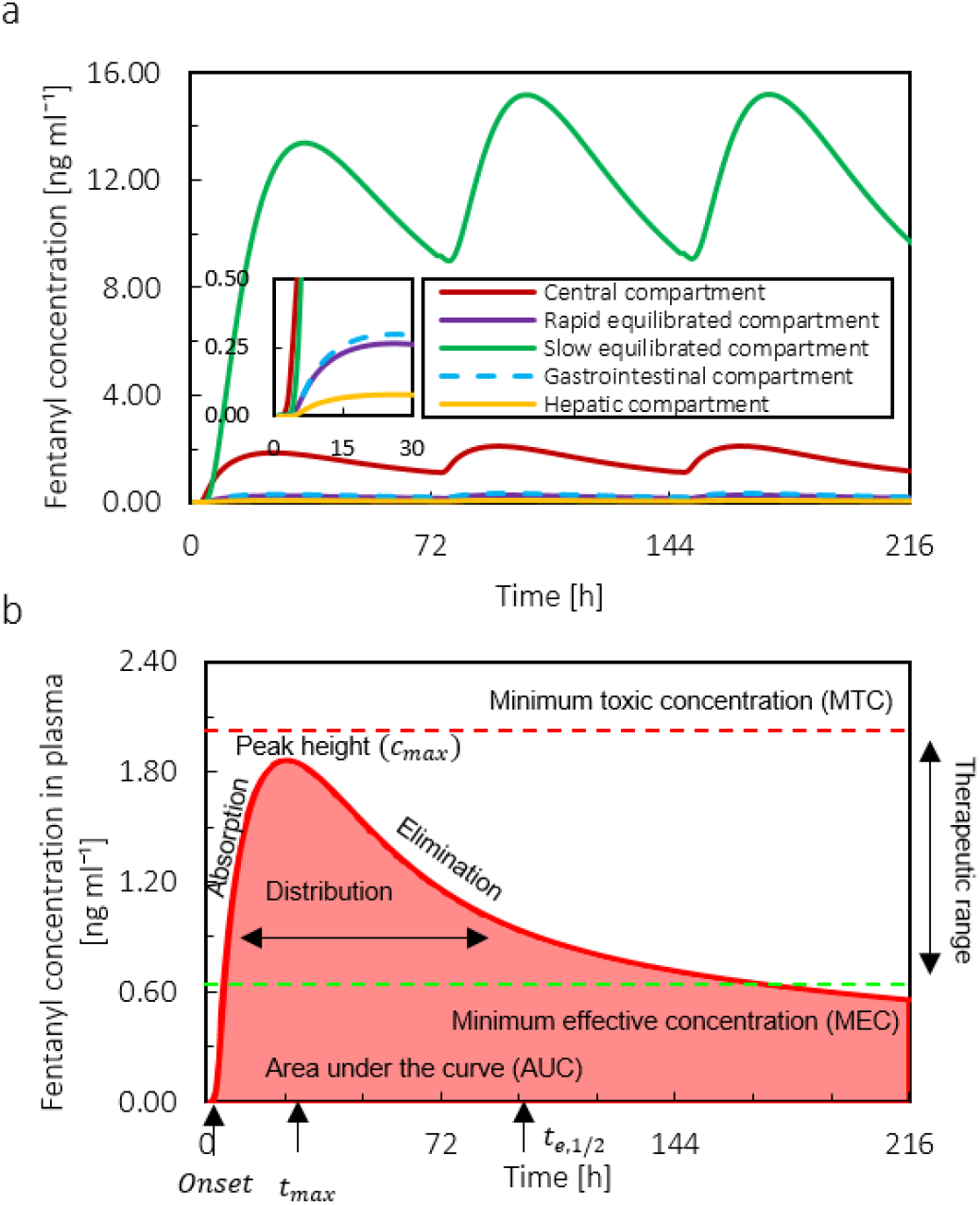
a: Fentanyl concentration in central, slow equilibrated, fast equilibrated, gastrointestinal, and hepatic compartment during 9 days by applying 3 patches each for 72 hours. b: Plasma fentanyl concentration during 9 days for applying only 1 patch (Duragesic® with the nominal flux of 75 µg h^-1^) for the virtual patient at the age of 20 years.

**Figure 8.**
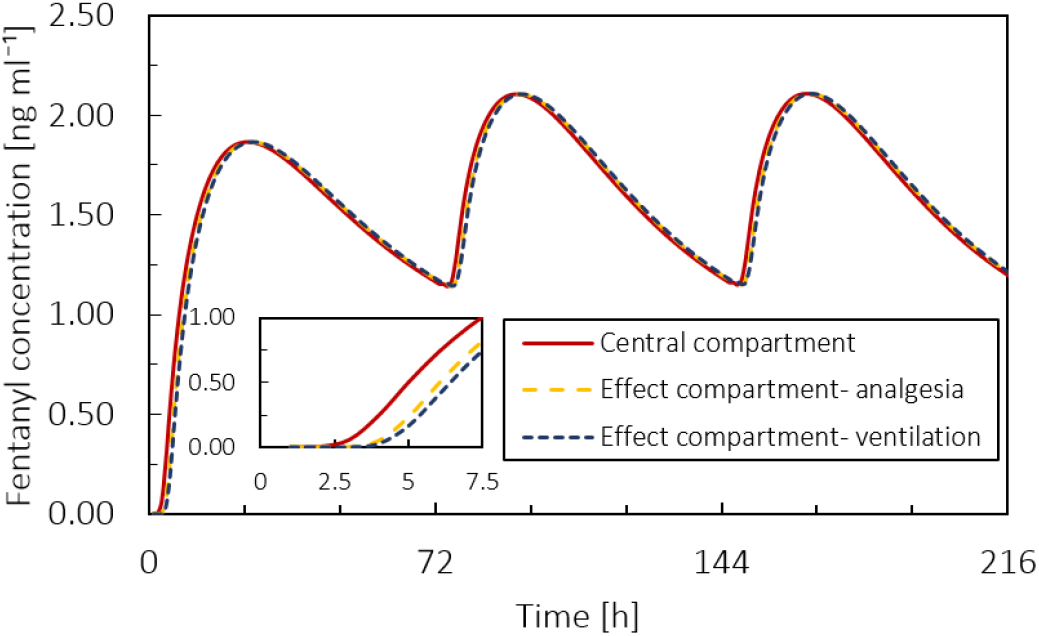
Fentanyl concentration in the central compartment, effect compartment for ventilation, and effect compartment for analgesia for 9 days by applying 3 patches each for 72 hours.

The initial amount of fentanyl in the patch was 12.6 *mg* for Duragesic® fentanyl patch with a target nominal flux of 75 µg h^-1^ (with 30 cm^-2^ surface area). After applying the patch, the drug diffuses from patch to skin and is eventually uptaken by the blood circulation system in the dermis. Figure 6/a shows that after 72 hours, the amount of fentanyl in the patch reduces from 12.6 mg to 8.99 mg, which means during 72 hours, only 28.6% of the drug in the patch was used. Simultaneously, the amount of fentanyl in the skin after 15 hours increased from 0 mg to reach its maximum at 0.843 mg, and then it decreased to 0.528 mg at the end of 72 hours. The difference between the total drug in the skin and patch together from the patch’s initial drug content is the amount of drug taken up from the skin by blood circulation. Figure 6/b shows that the fentanyl concentration in skin layers is shown, which shows that the fentanyl is more accumulated in the upper part of the skin close to the patch, as expected.

From the PK simulation, the concentration of fentanyl in each compartment for the virtual patient at the age of 20 years is shown in Figure 7/a. The concentration profile of fentanyl is based on the parameters mentioned in Table 2. The maximum plasma concentration for fentanyl after applying the first patch is 1.86 ng ml^-1^, which occurs 22 h after applying the first patch. Based on the literature, for Duragesic® fentanyl patch with the surface area of 30 cm^2^ the nominal flux is 75 µg h^-1^ [29]; however, based on our result, the fentanyl flux from the patch is highly time-dependent and far from being constant. The mean *C*_*max*_ of fentanyl is reported 1.7 (SD=0.7) ng ml^-1^ and the average time to reach the maximum concentration of drug in plasma is 33.5 h (SD=14.5) [29]. The difference between the obtained results from our simulation and reported averages are less than one standard deviation. The maximum fentanyl concentration in slow equilibrated, rapid equilibrated, gastrointestinal, and hepatic compartment is 13.96, 0.27, 0.31, and 0.08 ng ml^-1^, respectively. Time for maximum concentration (*t*_*max*_)for slow equilibrated compartment is 31 [h], and for the other compartments is 23 h. By comparison the concentration profile of fentanyl in slow equilibrated and central compartment, we realize due to lipophilicity of fentanyl, most of the drug is accumulated in adipose tissue. In Figure 7/b, the plasma fentanyl concentration (fentanyl concentration in the central compartment) for one transdermal patch over 216 h is shown. Based on the literature, the minimum therapeutic level for fentanyl is 0.63 ng ml^-1^, while the minimum toxic concentration is 2 ng ml^-1^ [46], [66]. For our base case, after 7.62 h, the fentanyl concentration in the virtual patient reached the therapeutic range (0.63-2 ng ml^-1^) and remained in this range for the following 208 hours. It should be mentioned that the therapeutic range is different for each person, and the shown therapeutic range is just an average.

In Figure 8, the fentanyl concentration in plasma is shown for the effect compartment of ventilation and analgesia. The calculation of concentration in effect compartment based plasma concentration is described in Equation 12 in section 2.1.3. The effect compartment only produces a delay between the concentration of fentanyl in plasma and the effect of the drug. As the result shows, the concentration profiles are very close to each other. This is due to the high potency of fentanyl, which facilitates the passage through the blood-brain barrier (BBB) [67].

Pain relief and reduction of ventilation rate for the patient after applying fentanyl patch are related to the concentration of fentanyl in the plasma. Based on the conventional therapy results for the virtual patient at the age of 20 years, as the plasma fentanyl concentration increased, the ventilation rate and VAS pain score dropped. When the fentanyl concentration in the plasma reaches its maximum, the rate of change in the plasma concentration decreases. The reduction in plasma concentration gradient also changes the gradient in pain intensity and reduction of ventilation rate. Therefore, after reaching the *c*_*max*_ of the first patch, the fluctuation in pain intensity and ventilation rate reduces.

### 3.2 Effect of age on the therapy

The stratum corneum is the main obstacle in the drug penetration path through the skin into the blood circulation system. As age increases, the thickness of the SC increases, and the thickness of the dermis decreases. Therefore, the increase in SC thickness will reduce the amount of fentanyl delivered to the body and lead to a delay and reduction of drug uptake. As shown in Figure 10/a, the effect of increasing thickness of the SC on drug flux out of the skin is more than the effect of decreasing thickness of the dermis. Based on this result, as age increased, the maximum flux decreased, and the time to reach this maximum flux increased. From age 20 to 80 years, the maximum flux of drug into the blood decreased by 11%, and the time to reach this maximum flux increased by 15%. This implies that by applying the same fentanyl patch for the patient at different ages, the patient uptakes less amount of fentanyl by aging. Simultaneously, the time to reach the maximum flux of the drug increases.

**Figure 9.**
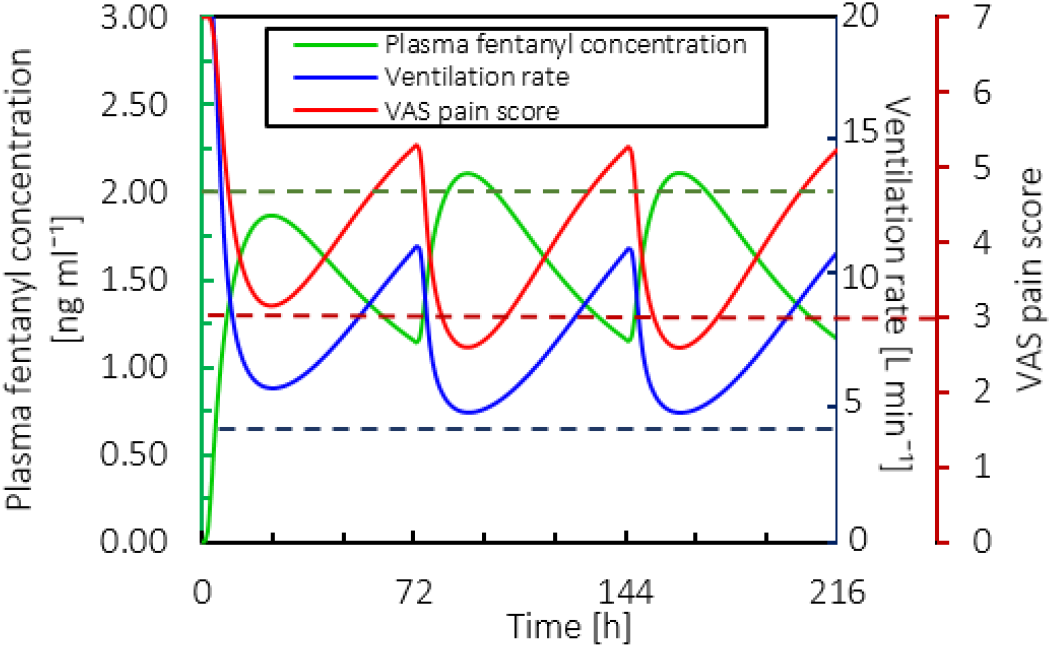
Ventilation rate, VAS pain score, and plasma fentanyl concentration for the base case for 9 days by applying 3 patches each for 72 hours.

**Figure 10.**
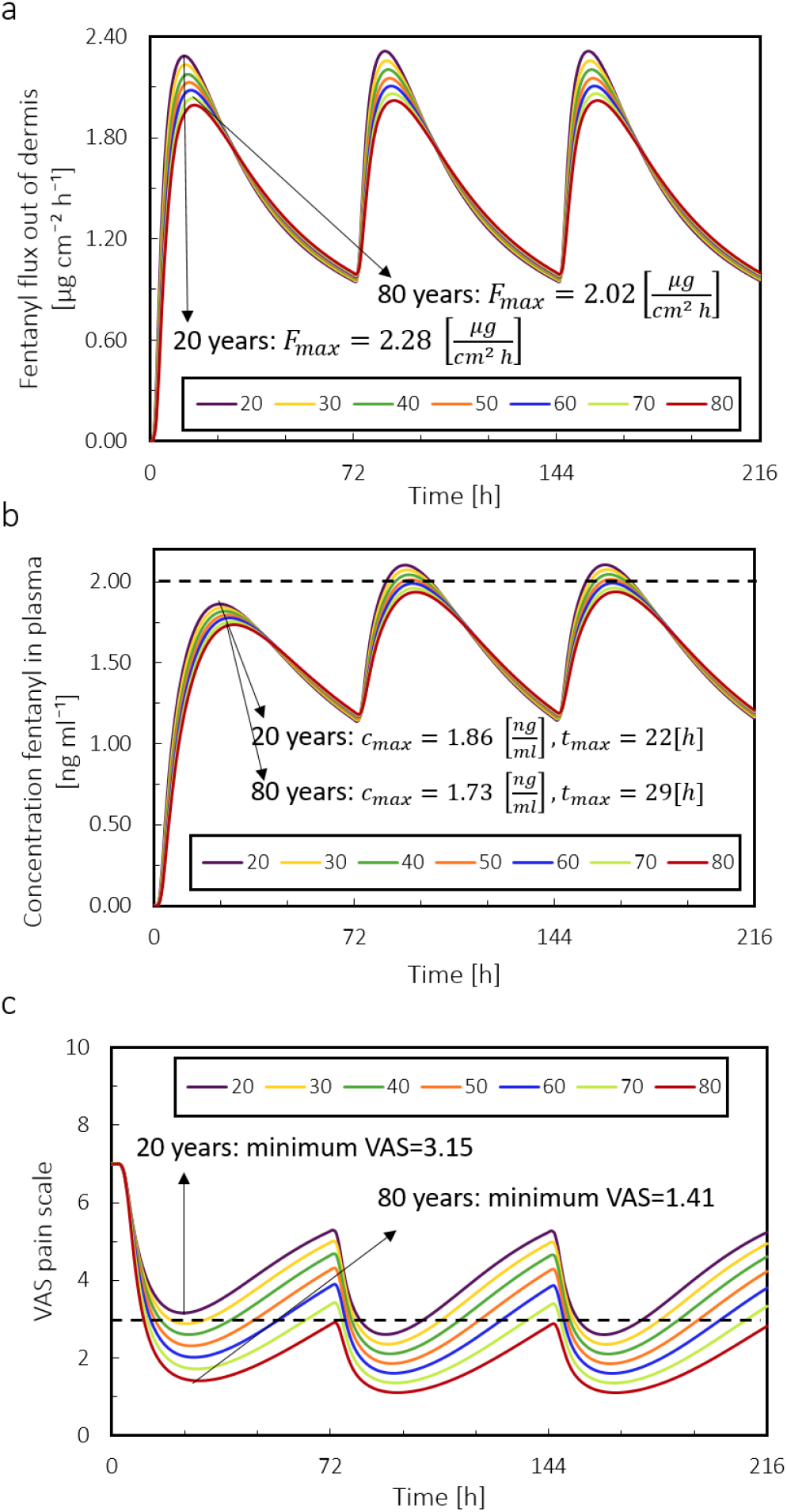
a: Fentanyl flux out of dermis; b: Plasma fentanyl concentration. The dotted line is concentration threshold in plasma; c: VAS pain score for the virtual patient with different ages from 20 years old to 80 years old during 9 days by applying 3 patches each for 72 hours. The dotted line is the pain intensity target.

In this study, we assumed that the PK parameters do not change with age. However, as the age changes, the drug flux out of the dermis will change. Consequently, the different blood drug uptake, which is a result of different drug flux out of the dermis, changes the concentration of drug in PK compartments. Figure 10/b shows that the maximum concentration of fentanyl from age 20 to 80 years decreases by 7%, and the time to reach this maximum increased by 32%. The reduction of *c*_max_ and the increase in t_max_ by increasing age are also reported in other works such as Thompson et al. (1998) and Paut et al. (2000). The result implies that even if the patient at different ages receives the same therapy, by aging, the accessible drug amount for the patient for pain relief (concentration of drug in plasma) reduces.

Based on the result of drug uptake and PK modeling, the virtual patient at older ages received less amount of drug, which led to lower fentanyl plasma concentration. As mentioned in the material and method section (section 2.4.2), the fentanyl concentration required to reach the half-maximum effect will be reduced by aging. The result of PD, shown in Figure 10/c, shows that, despite receiving less fentanyl, the VAS pain score in older age was reduced more than in younger age. The reduction in VAS pain score for the patient at the age of 80 years was 55% more than the age of 20 years. The reduction of required opioids for pain relief with increasing age is reported in other works too [65], [70]–[72]. To summarize the result of this section, if a similar fentanyl therapy is being applied for the patient, by increasing the age, the patient receives less amount of drug. However, the patient might still experience more pain relief as opioid requirements for pain relief decreases by age.

### 3.3 Individualized therapy with a precalibrated digital twin

As mentioned in the previous section, virtual patients of different ages received different amounts of the drug and showed different levels of pain relief. Therefore, it is actually needed for each virtual patient in a certain age category to receive a different therapy to reach desirable pain relief. Using a precalibrated digital twin of patients in a certain age category, we composed a more suitable therapy for each age category. The fentanyl concentration and VAS pain score for different age categories are shown in Figure 11/a and b. The results in these two graphs show that the precalibrated therapy by digital-twin successfully kept the pain intensit below the target by slightly increasing in concentration of fentanyl in plasma and keeping it in almost constants value. This increase in concentration was only done by changing the frequency of replacing the patch. Another criterion in the proposed therapy by digital twin was avoiding respiratory depression, which we monitored by ventilation rate. Results in Figure 11/c show, despite the increase in the concentration of fentanyl in plasma, the digital-twin suggested therapy tried to keep this increasing at a minimum level to avoid hypoventilation.

**Figure 11.**
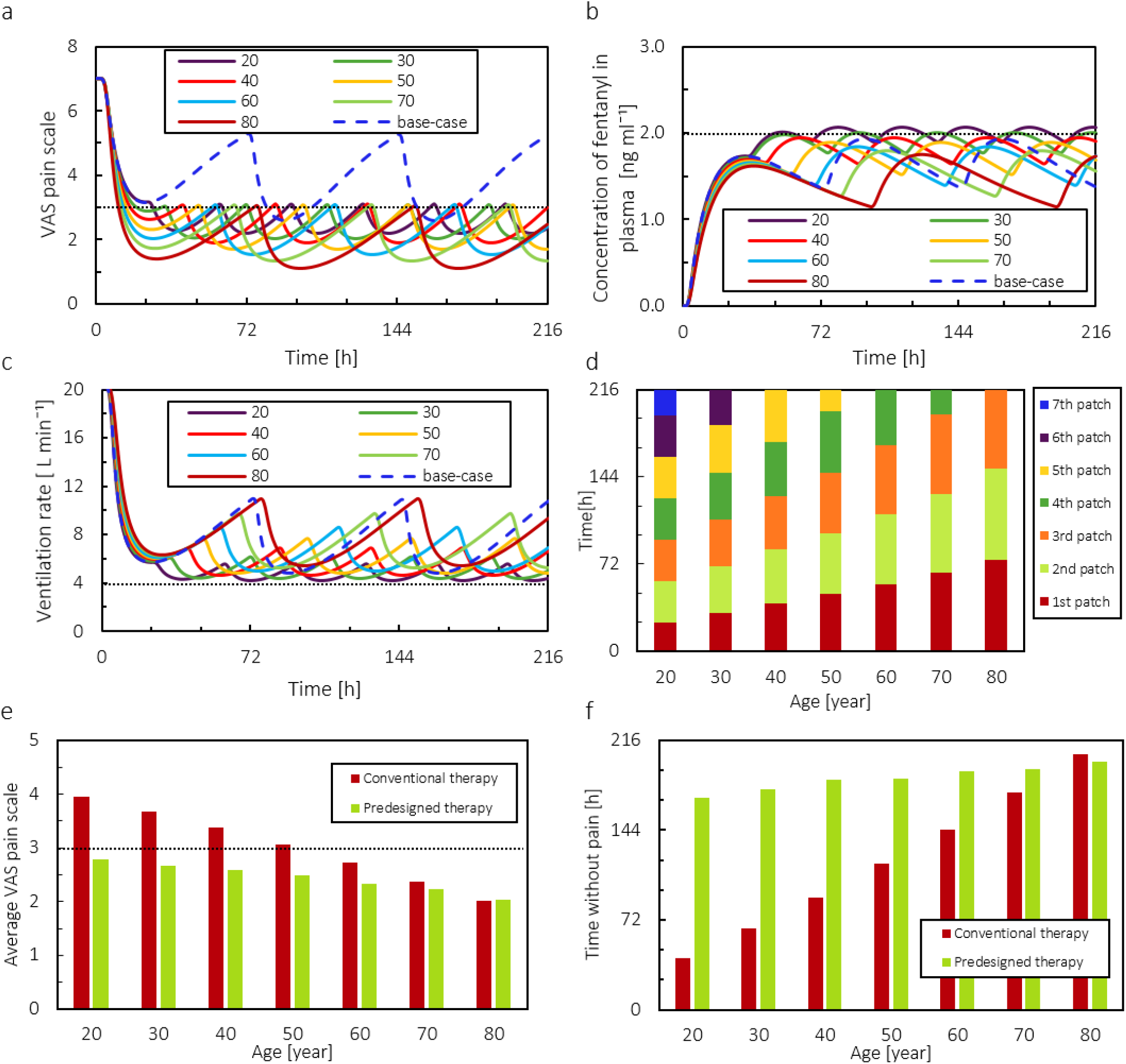
a: VAS pain score versus time for predesigned therapy. The dotted line represents the target pain intensity. The base case is conventional therapy for 20 years old virtual patient. b: Fentanyl plasma concentration versus time for predesigned therapy. The dotted line represents the threshold for fentanyl concentration in plasma. The base case is conventional therapy for 20 years old virtual patient. c: The ventilation rate versus time for predesigned therapy. The base case is conventional therapy for 20 years old virtual patient. d: Duration of applying each patch. The dotted line represents the minimum normal ventilation rate. e: The average VAS pain score both for conventional therapy and for proposed pain control therapy. The dotted line represents the target pain intensity. f: The time that patient VAS pain score was under 3, which we assumed as time without pain for virtual patients with 20 to 80 years old during 9 days both for conventional therapy and proposed pain control therapy.

As shown in Figure 11/d, the virtual patient at different ages needs a different number of patches in the same time frame, with different durations of application. In conventional 3-day therapy for 216 hours, only 3 patches will be applied. However, for achieving a therapy where the pain score shall be kept under 3 for the age of 20 years, almost 7 patches are needed. Based on these results, by aging, the patient needs to change the patch less frequently. The result in Figure 11/e shows that by applying the proposed age-dependent therapy, the average pain intensity for the patient at the age of 20 years decreased by 30%. Therefore, the change in time of replacing the patch led to better performance of proposed therapy by digital twin compare to conventional therapy in relieving the pain. Note that we could also adjust and change therapy by using patches with different concentrations as well, but this we did not explore.

As mentioned in previous paragraphs, the aim of the proposed therapy by digital twin was to keep the VAS pain score under 3 while keeping the plasma concentration at the lowest possible amount to reach this target. Here we calculated the duration of which the pain intensity was below 3, which we call time without pain, as a parameter to evaluate the success of the treatment. The result in Figure 11/f shows that the digital twin increased the time without pain considerably. For the virtual patient at the age of 20 years, the time without pain increased by 314%. As mentioned in section 2.4.3, decreasing pain intensity to a much lower value than 3 is not the aim of the therapy; however, it is important to keep it under 3 while avoiding therapy’s adverse effects. By implementing the digital twin, we were successful in keeping the VAS pain score under 3 for a longer period of time. Therefore by considering the presented result in Figure 11, the digital twin successfully decreased the pain intensity and increased the time without pain. Meanwhile, the increase in the concentration of drug in plasma was at a controllable level, in which the virtual patient did not face hypoventilation in the period of the therapy. The fluctuations in drug concentration, VAS pain score, and ventilation rate were the smallest for a young patient as they need to replace the patch more frequently.

The predesigned therapy by a precalibrated digital twin proposed that 3 patches with the nominal flux of 75 µg h^-1^ for 9 days are not sufficient in order to reach the favorable pain relief for the patient at the age of 20 years. However, it would suffice for an 80-year-old patient. This therapy suggested the patch needs to be changed more frequently to keep the VAS pain score under 3. In the clinics, when the applied fentanyl patch is not efficient for pain relief, there are two other options. First, change the patch at each 48 h instead of 72 h [13]; second, change the patch to a higher dose. Here we compared the performance of four different therapies for the virtual patient at the age of 20 years: 1. Predesigned therapy by digital twin with fentanyl patch with the nominal flux of 75 µg h^-1^; 2. Using fentanyl patch with the nominal flux of 100 µg h^-1^ and changing at each 72 h so conventional therapy at a higher dose; 3. Using fentanyl patch with the nominal flux of 75 µg h^-1^ and changing at each 72 h so conventional therapy; 4. Using fentanyl patch with the nominal flux of 75 µg h^-1^ and changing at each 48 h. It should be noted the last three therapies in the list are conventional therapies that are used in the clinics.

The result of this comparison is shown in Figure 12. The maximum concentrations of fentanyl in plasma for all these four approaches are above 2 ng ml^-1^ threshold, but it was necessary to reach the target pain relief (Figure 12/a). The result shows by digital twin therapy, the VAS pain score is at the favorable level (under 3) and time without pain has the highest value compared to the other three approaches (Figure 12/b and c). The second approach (using fentanyl patch with the nominal flux of 100 µg h^-1^ was not successful in keeping the ventilation rate in the normal range (Figure 12/d and e), while the other three approaches managed to do it. However, it was successful in keeping the VAS pain score below 3. As in predesigned therapy by digital twin, a change of the patch (with the nominal flux of 75 µg h^-1^) is needed more frequently; the fraction of unused drug in the patch for this approach compared to the other three approaches is higher by 17.1%, 17%, and 9.3% respectively (Figure 12/f). To summarize this comparison, using the patch with the nominal flux of 75 µg h^-1^ and changing every 72 h (third approach) or using the patch with the nominal flux of 75 µg h^-1^ and changing every 48 h (fourth approach) is not very effective in pain relief. Therefore, based on the main criteria for pain management, the best solution can be chosen. For the aim of keeping the pain under VAS pain score 3, the first and second approaches can meet the requirements. On the other hand, the second approach was not able to avoid hypoventilation. When we want to achieve sufficient pain relief (VAS < 3) and a sufficiently high ventilation rate, only predesigned therapy by digital twin meets all the criteria. However, it should be noted the frequency of changing the patch affects the cost of the therapy as more patches are needed.

**Figure 12:**
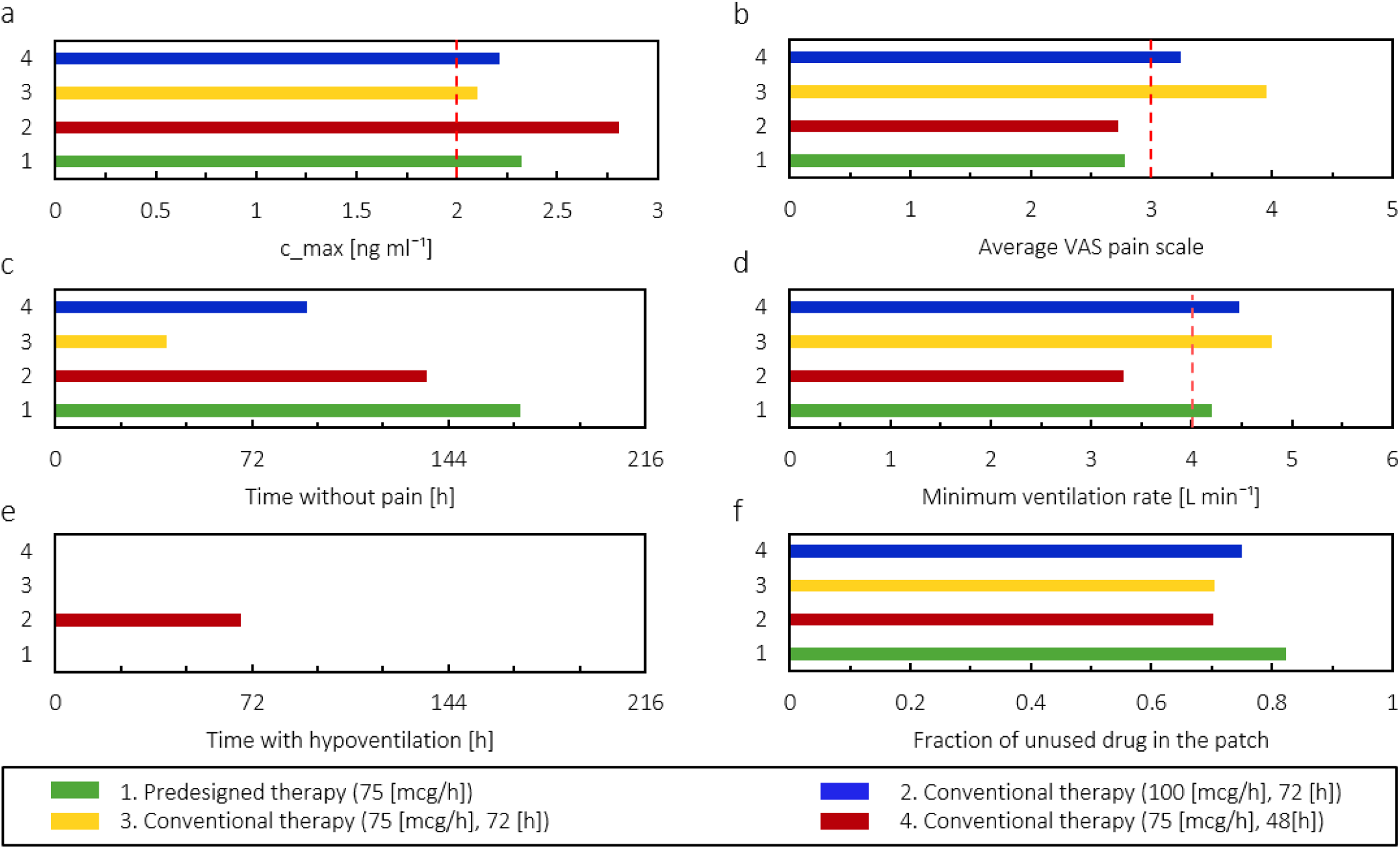
comparison of four therapies (predesigned therapy with digital twin and three conventional alternative therapies). a: maximum concentration of fentanyl; b: average VAS pains score, c: time without pain; d: minimum ventilation rate, e: duration of hypoventilation; f: the fraction of unused drug in patch during 9 days for the virtual patient at the age of 20 years.

### 3.4 Individualized therapy by the real-time digital twin

To evaluate the effect of patient feedback in the digital twin’s proposed therapy, we assumed 100 virtual patients with the same physiological state as the 20 years virtual patient. This means all these 100 virtual patients had the same model input parameters but varied in their feedback response to the treatment. These virtual patients fed their VAS pain score into the model based on the procedure mention in section 2.4.4. In this approach, sometimes, the patient will thereby not receive the required amount of fentanyl, and sometimes, the patient receives an extra amount of drug, which is not necessary. As a result of including patient feedback, the pattern of replacing the patch was changed in order to meet the patient’s needs. To compare the performance of this feedback approach with the one in section 3.3, we evaluated the average fentanyl plasma concentration, VAS pain score, time without pain, and ventilation rate.

In Figure 13/a, the result shows that in most cases, the average fentanyl concentration was lower compare to the therapy proposed by the precalibrated digital twin. If we apply a t-test on the result of these two therapies, the obtained p-value is 1.8 x 10-8, which implies the difference in the plasma concentration for both approaches is significant. The blue boxplot at each graph represents the difference between the corresponding values between two approaches for every single patient. In Figure 13/b, the average VAS pain score via the two approaches is shown. Based on the result, the median of average VAS pain score is higher for the therapy with the feedback. However, the p-value for pain intensity between these two approaches is 0.32, which implies there is no significant difference in the average VAS pain score via these two approaches. Therefore, we can conclude including patient feedback does not have a significant effect on the pain relief performance of the digital twin.

**Figure 13.**
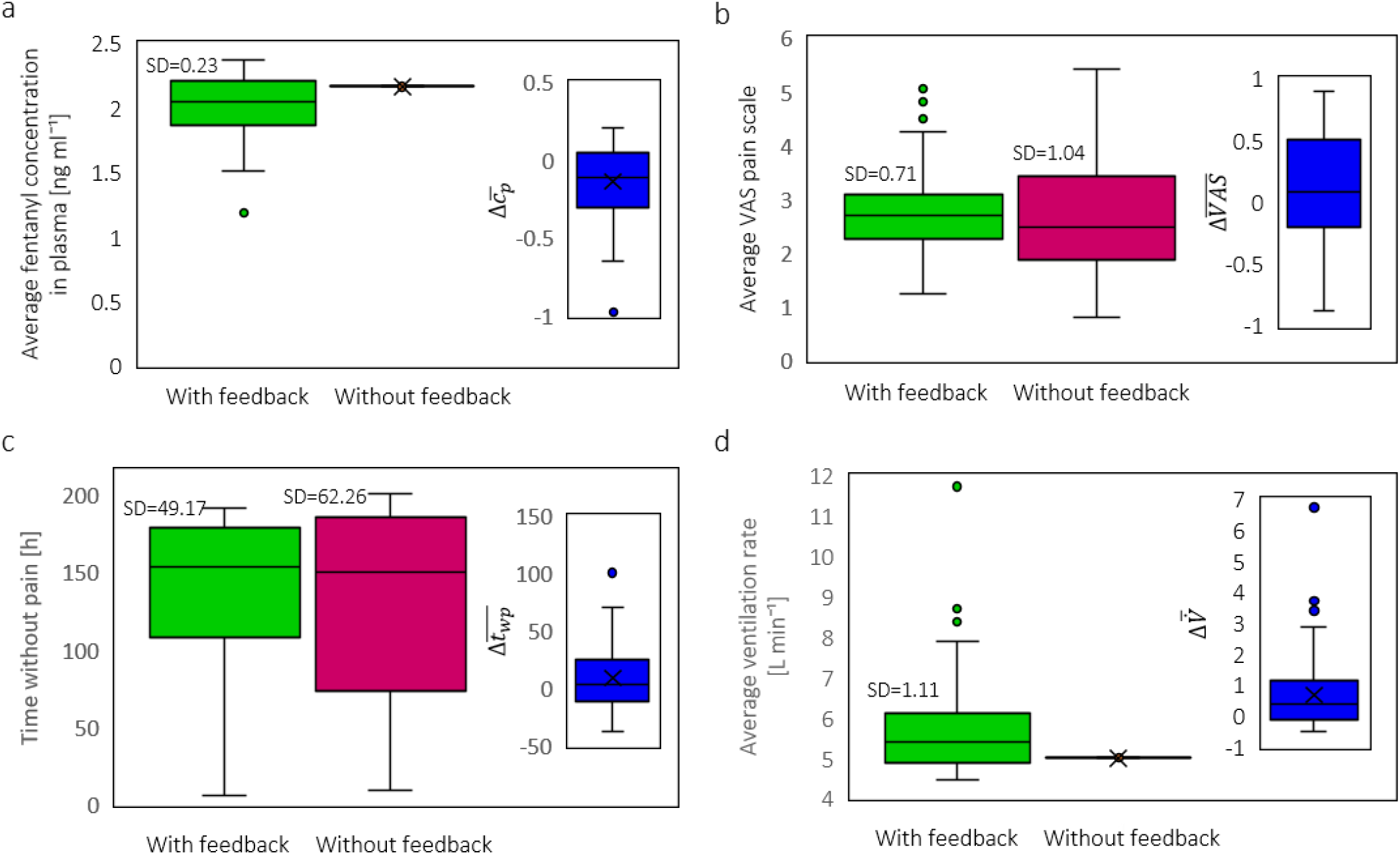
a: Distribution of average plasma concentration of fentanyl; b: Distribution of average VAS pain score; c: Distribution of time without pain; d: Distribution of average ventilation rate for the same population of 100 virtual 20 years old patients for 9 days both for pain control therapy by including feedback and without including feedback. The results are for the same virtual population with 100 members.

In Figure 13/c, the time without pain for each case via two approaches is shown. The results show that the median average VAS pain score is higher by 3 hours for the digital twin with patient feedback. Additionally, the standard deviation for the digital twin with the patient feedback approach is lower than without the patient feedback approach by 10%. However, the p-value of the difference between times without pain via these two approaches is 0.25, which implies no significant difference between them. In the digital twin with patient feedback approach, when the patch is changed is based on the updated VAS pain score, the fentanyl plasma concentration varies individually. Therefore, the ventilation rate will be different for all these 100 cases. In the digital twin without patient feedback approach, as the patches will change at the same time for all these 100 cases, the plasma fentanyl concentration will be the same for all of them; therefore, the average ventilation rate was 3.9 L min^-1^ for all the cases. The result in Figure 13/d demonstrates that by considering feedback, more than 50% of the patients have a higher ventilation rate than the approach that we did not consider the patient’s feedback. This is the result of not delivering an extra amount of drug to the patient. For the patients with a lower ventilation rate, the patient needed more amount of fentanyl to reduce the pain level. Therefore, the ventilation rate was lowered. The decision-making criteria to change the patch in this study were done only based on the VAS pain score, not a combination of VAS pain score and ventilation rate. The result of the t-test for ventilation rate via these two approaches is 5.9 x 10-8, which refers to a significant difference between these two approaches. We can conclude that the reduction in plasma concentration and an increase in ventilation rate makes the real-time digital twin therapy safer for the patient. While including patient feedback in the developed digital twin did not significantly change the digital twin’s performance in pain relief.

## 4. OUTLOOK

In this study, several model simplifications were considered for drug uptake, pharmacokinetics, and pharmacodynamics modeling. Not all processes that happen in the human body are exactly captured as in reality. Nevertheless, the models were validated and showed a good agreement with experimental data. Besides these assumptions, it is important to develop the digital twin based on real humans and connect the digital twin to the real patient by real-time sensor data or patient feedback. In order to obtain a more accurate digital twin to assist in the treatment, we should consider other factors. Here are some of the important model improvements that should be applied:

- In this study, the skin’s geometry was a simple layered structure with no heterogeneity, and perfect contact between the patch and the surface of the skin was assumed. Developing a real structure of the skin by considering its geometry and different diffusion paths and skin components will lead to a more accurate drug flux from the skin. However, it should be considered that the skin structure is very complex and differs depending on its location and the individual. If the modeling goes to a high level of individualization, every person and location must be measured, as literature data will be invalid. This may create new problems with the relevance and reliability of the measurement method. Therefore, a balance has to be found between the individualization/realism of the model and its practicability in order not to create new model artifacts that are greater than for a simplified model due to unreliable measurements of individual skin parameters.
- Advection - which is an important mechanism in drug penetration - could be included by considering the dermis’ capillary vessels. In this study, the capillary network was not considered explicitly but accounted for in the model through an equivalent diffusion length of the dermis.
- As mentioned in the sensitivity analysis section (Supplementary material), the fraction of unbound drug has an important role in the plasma’s calculated fentanyl concentration. In our model, a constant value was considered based on literature. Considering its reaction’s kinetics would lead to different values as a function of time and concentration for each patient, which will increase the accuracy of the digital twin.
- An important issue in opioid therapy is the changing opioid tolerance of the patient through the use of opioids over time. Increasing this tolerance will increase the required drug for pain relief; therefore, it could be considered in the model.
- Here we only controlled therapy, so the changing of the patches, based on the pain level of the patient. Reducing the adverse effects is crucial to reach an effective and safe treatment; it is important to control therapy based on therapeutic effects and adverse effects.
- Besides the impact of age between different patients, also other factors need to be considered. Physiological features of the patients, such as gender, weight, or background disease, can play an important role in the outcome of fentanyl therapy.
- To analyze the digital twin’s performance in providing efficient and safe treatment, it is important to develop a digital twin based on the real patient. In this case, the twin can be updated with patient feedback over time. The result of the digital twin precalibrated therapy can then be compared with the real result to validate each individual patient’s digital twin’s performance. It should be noted that the assessment of the parameters of the real patient may introduce new errors due to the accuracy and reliability of the used sensors.

## 5. CONCLUSIONS

In this study, we developed a physics-based digital twin for fentanyl transdermal therapy. With this twin, we first anticipated the outcome of conventional transdermal therapy on a patient of different ages. In addition, based on the age of the patient and the corresponding response of the patient to treatment, the digital twin was used to propose a predefined alternative transdermal therapy for the patient at each age, namely by changing the patches at a different time interval. Finally, the proposed therapy by digital twin was updated by virtual patient feedback to have a more accurate and safer therapy.

Based on the result of the simulation by aging, the patient will receive less amount of drug as a result of the increase in the thickness of the stratum corneum for conventional therapy that is now used in the clinics. However, as the required amount of drug for pain relief decreases by age, older patients still experienced more pain relief, despite a lower blood concentration. By applying conventional therapy, younger patients had a maximum fentanyl flux out of the dermis that was 11% higher, the maximum concentration of fentanyl that was 7% higher, and pain relief that was 55% lower than older patients. When the digital twin proposed a different therapy for each age category, younger patients need to apply patches more frequently compared to the older patient to reach the target pain relief. This twin-assisted therapy reduced the VAS pain score by 30% and time without pain increased by 314% for a patient at the age of 20 years, compared to conventional therapy. In the next step, the digital twin was included patient feedback on their pain intensity at certain times. With such feedback, the therapy was better tailored to real-time patients’ needs and avoided delivering insufficient or extra fentanyl. By including the patient’s feedback in the digital twin, we successfully were able to avoid hypoventilation, which means the therapy was safer for the patient.

The proposed physics-based digital twin was built up based on a real patient’s physiological features. These features are age, body mass, gender, and other involved factors that play an important role in fentanyl pharmacology. Real-time patient feedback for pain relief and oxygen saturation levels in the blood will improve the therapies proposed by the digital twin. The proposed therapies by a digital twin for each patient can increase the concentration of the drug at a level to reach the required pain relief while avoiding its adverse effects. We quantified the added value of a patient’s physics-based digital twins and sketched the future roadmap for implementing such twin-assisted treatment into the clinics.

## Acknowledgments

This work was financially supported by the Novartis Research Foundation (grant “Virtual twinning for intelligent, personalized transdermal drug delivery”). The funder was not involved in the study design, collection, analysis, and interpretation of data, the writing of this article, or the decision to submit it for publication.

## Abbreviation

NSAIDs: Non-steroidal anti-inflammatory drugs
PBPK: Physiologically-based pharmacokinetics
SC: Stratum corneum
TDDs: Transdermal drug delivery systems
VAS: Visual analog scale

## Notes

### Competing Interest Statement

The authors have declared no competing interest.

