## supplementary material for "Predicting transdermal fentanyl delivery using physics-based simulations for tailored therapy"

### S.1. Sensitivity Analysis over the parameters

The results of the sensitivity analysis over model input parameters are demonstrated in Figure s1. In Figure s1/a, the sensitivity index of the fentanyl flux out of the dermis to input parameters for the drug uptake model is shown. These input parameters are partition and diffusion coefficient in the patch and skin layers and the thickness of the skin layers thickness. Based on this result, the fentanyl flux was most sensitive to epidermis thickness and diffusion in the epidermis. Based on the sensitivity index, the flux is not so sensitive to equivalent dermis thickness; however, this result is for a 1% change in the value of the equivalent dermis thickness. It should be noted that the changes in equivalent dermis thickness could be notable. Its thickness can change from not considering the dermis layer (assuming drug uptake at the interface of the epidermis and dermis) to considering the whole thickness of the dermis (a few millimeters). Therefore, despite the obtained low sensitivity index for dermis thickness, dermis thickness could play an important role in the outcome of the model.

In Figure s1/b, the sensitivity of the average plasma concentration to the volume of the compartments, inter-compartmental clearance, renal and hepatic clearance, and the fraction of unbound drug is shown. The average concentration of fentanyl in the plasma is largely sensitive to the fraction of unbound drug. Between the blood and other compartments, and the only unbounded drug can be transferred; therefore, this parameter has a huge impact on fentanyl concentration in plasma. Another important parameter in the PK model is the blood flow to the gastrointestinal compartment. Unlike other compartments, in which outgoing flow gets back to the central compartment, for the gastrointestinal compartment, the outgoing flow goes to the liver. Therefore, the blood flow of the gastrointestinal compartment will affect the amount of metabolized the drug. The sensitivity index of the average VAS pain score to the Half maximal effective concentration and Hill coefficient in Figure s1/c shows that the concentration of half-maximum effect has an important role in the resulting effect.

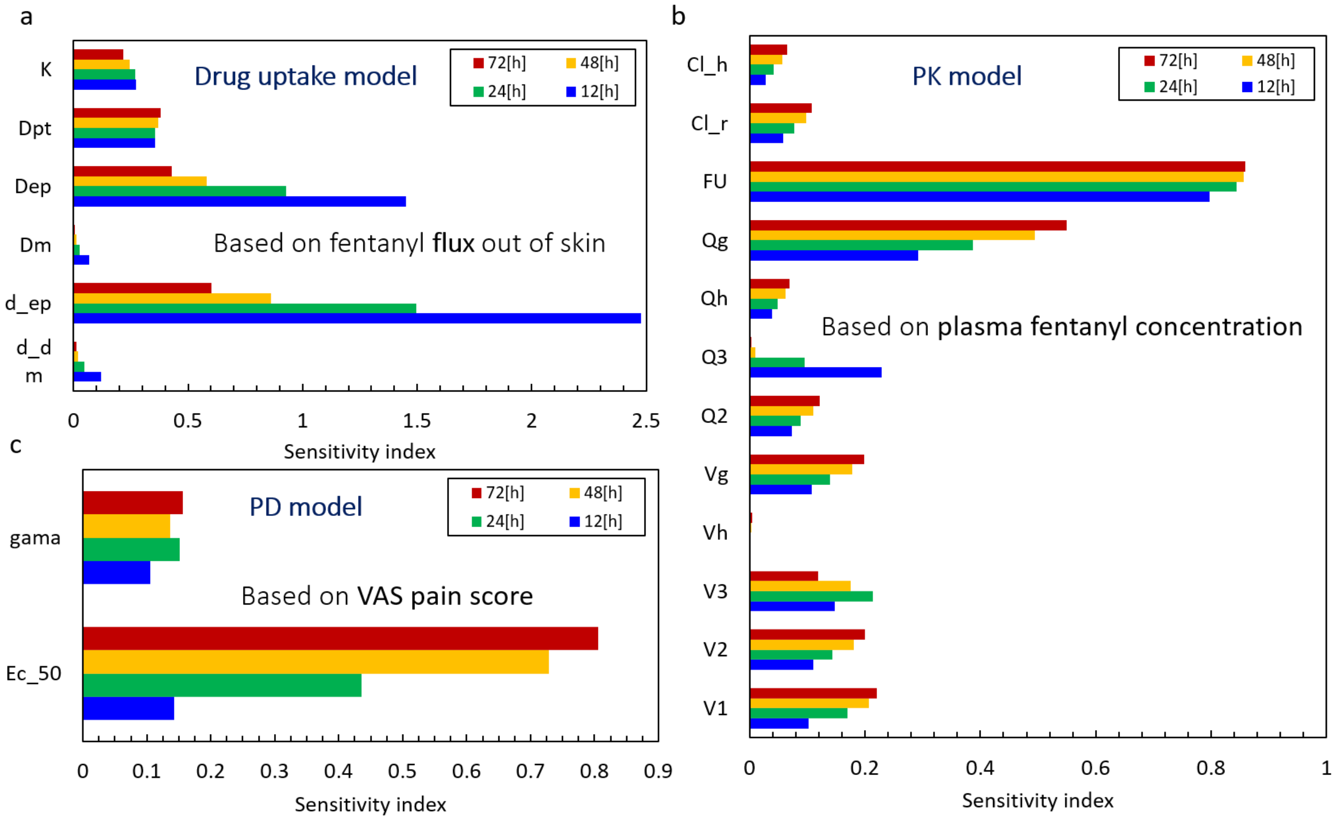


***Figure s1- Sensitivity analysis results, a: sensitivity index of fentanyl flux out of dermis to drug uptake model parameters. b: sensitivity index of plasma fentanyl concentration to PK model parameters. c: sensitivity index of VAS pain score to the PD model parameter. The analysis was done over 72 hours of therapy for the virtual patient at the age of 20 years with Duragesic® fentanyl patch with the nominal flux of 75 µg $h^{-1}$***

### S.2. Validation of the PK model

To evaluate the validation of results from the pharmacokinetic model, we compared our result with Marier et al. (2006), in which the detail of the experiment is provided in section 2.1.4.2. The results of the simulation and the average results of the experiments [6] are shown in Figure 5. The root-mean-square deviation (RMSD) of simulation data and experimental data was 0.152 [ng $ml^{-1}$]. By analyzing the result from simulation and the experiment, we find at the beginning there is a time difference for reaching the maximum concentration for the first peak; however, this difference is less for the next two peaks. In the experiment, in the third peak, there is a jump in the concentration, which the simulation cannot predict. This jump could be due to changes in the situation for patients or the measuring process of concentration of fentanyl in plasma. However, in general, the agreement of the model with the experiments is satisfactory.

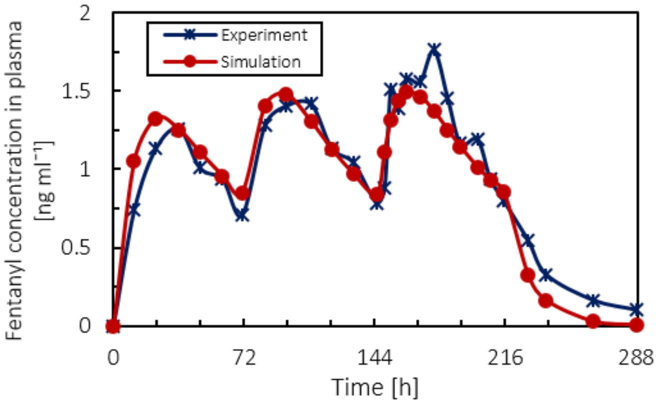


***Figure s2- Concentration of fentanyl in the blood circulation system ($c_p$) as a function of time from experiment and digital twin over 11 days by applying 3 patches with the nominal flux of 50 µg $h^{-1}$, each for 72 hours following two days with no patches.***

### S.3. Calibrating the PD model for VAS pain score

As mentioned in section 2.1.4.3, we obtained the value of constants for sigmoid function related to the relationship between VAS pain score and fentanyl concentration in the effect compartment (Equation 21) based on experimental data. In Figure 6, the average experimental VAS pain score [62] and simulated PD model data are shown. The corresponding simulated plasma fentanyl concentration is demonstrated in the subplot. Based on the fitted sigmoidal model, as concentration increases, the pain intensity decreases. For instance, from t=4 h to 8 h the patient's pain intensity has a plateau while the concentration of fentanyl is increasing in the blood and subsequently in the central nervous system. This different behavior could be due to several reasons such as the time lag between drug concentration and effect, deviance from measure concentration, and the overall average concentration of drug in plasma. However, in general, the agreement of the model with the experiments is satisfactory, and the percentage error is within 8.9%.

$$E_{VAS} = E_{0,VAS} - E_{max,VAS} \times \frac{c_{e,VAS}^{\gamma}}{EC_{50,VAS}^{\gamma} + c_{e,VAS}^{\gamma}} \tag{S1}$$

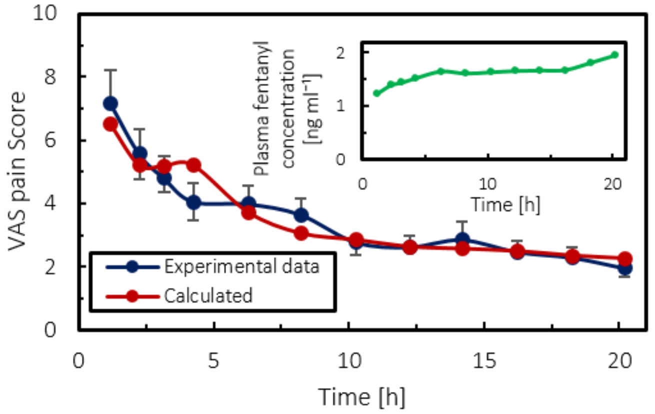


***Figure s3: Average experimental VAS pain score during fentanyl therapy and fitted data. Subplot: concentration of fentanyl in the blood circulation system, for infusion rate of 1.0 µg $kg^{-1}$ $h^{-1}$.***
